# Cytoskeletal regulation of a transcription factor by DNA mimicry

**DOI:** 10.1101/2021.09.30.462597

**Authors:** Farah Haque, Christian Freniere, Qiong Ye, Nandini Mani, Elizabeth M. Wilson-Kubalek, Pei-I Ku, Ronald A. Milligan, Radhika Subramanian

## Abstract

A long-established strategy for transcription regulation is the tethering of transcription factors to cellular membranes. In contrast, the principal effectors of Hedgehog signaling, the Gli transcription factors, are regulated by microtubules in the primary cilium and the cytoplasm. How Gli is tethered to microtubules remains unclear. We uncover DNA mimicry by the ciliary kinesin Kif7 as a mechanism for the recruitment of Gli to microtubules, revealing a new mode of tethering a DNA-binding protein to the cytoskeleton. Gli increases the Kif7-microtubule affinity and consequently modulates the localization of both proteins to microtubules and the cilium tip. Thus, the kinesin-microtubule system is not a passive Gli tether but a regulatable platform tuned by the kinesin-transcription factor interaction. We re-tooled the unique DNA-mimicry-based Gli-Kif7 interaction for inhibiting the nuclear and cilium localization of Gli. This strategy can be potentially exploited for downregulating erroneously activated Gli in human cancers.

## Main text

The tethering of transcription factors to membranes of cytoplasmic organelles (e.g. the endoplasmic reticulum, Golgi and mitochondria) or the plasma membrane, and their signal-dependent trafficking into the nucleus is a well-established mechanism for the precise regulation of gene repression and activation^1^. In recent years, the microtubule cytoskeleton has emerged as an important scaffold for tethering transcription factors away from the nucleus for pathway repression^2–4^. However, the molecular and structural mechanisms by which the cytoskeleton regulates transcription factors remains poorly understood.

An essential developmental pathway with a strict and conserved requirement for the microtubule cytoskeleton is Hedgehog (Hh) signaling^5–8^. In vertebrates, this pathway has a unique dependence on the primary cilium^9^, a microtubule-based organelle. Central to regulation of the vertebrate Hh signaling is the trafficking and accumulation of Gli proteins (Ci*; Drosophila Melanogaster* homolog) in the primary cilium. The Gli family of proteins consist of three isoforms Gli1, Gli2 and Gli3. Of these proteins, Gli1 is a transcriptional activator^10^, and Gli2 and Gli3 are dual-transcription factors characterized by the following domain organization: a repressor domain at the N-terminus and a trans-activator domain at the C-terminus that are separated by DNA-binding zinc-finger domain and sites for proteolytic cleavage^11, 12^. During Hh pathway repression, Gli2 and Gli3 are proteolytically processed into the shorter repressor form at the base of the cilium, with Gli3 acting as the major repressor^13–15^. Activation of the pathway results in increased accumulation of the full-length activator forms of the Gli proteins at the distal cilium tip, with Gli2 acting as the major activator, which promotes the expression of Gli1^16–19^. In vertebrates, defects in cilium assembly and architecture, result in incorrect processing of the Gli proteins^20, 21^. These observations suggest that the microtubule cytoskeleton plays an active role in Hh pathway activation. However, the mechanism of microtubule-dependent Gli regulation remains poorly understood.

The non-motile ciliary kinesin, Kif7 (ortholog of the *Drosophila melanogaster* Costal-2 kinesin-like protein and homolog of the motile ciliary kinesin Kif27), is a core Hh pathway component that provides the molecular link between the microtubule cytoskeleton and Gli regulation^22–25^. Loss of function mutations of Kif7 in zebrafish and mice embryos show defects in Hh-dependent developmental patterning^22–24, 26^, and clinical mutations in Kif7 have been associated with severe ciliopathies^27, 28^. When the Hh pathway is off, Kif7 is largely present at the base of the cilium, where it is thought to promote the formation of the repressor form of Gli. Upon pathway activation, both Kif7 and Gli accumulate at the tip of the primary cilium, a step required for Gli activation^25^. The precise mechanism for the localization of Kif7 to the base of the cilia is not known and multiple mechanisms have been suggested for its trafficking and cilium tip localization^25, 29–33^. In the absence of Kif7, Gli can enter the cilium but is not correctly concentrated at the distal tip of the axoneme^30^. These observations together with the reported co-precipitation of Gli proteins with Kif7 in cell and embryo lysates^23^ ^h^ave led to the proposal that microtubule-bound Kif7 establishes a ‘cilium-tip compartment’, a passive platform at the distal cilium tip, to which Gli is recruited^25, 30^. However, although current models assume a direct binding between Gli and Kif7, this has not been conclusively established and the molecular and structural basis of their interaction remains elusive. Furthermore, it is unknown whether the Kif7-Gli interaction is sufficient to tether the transcription factor to microtubules, and how the Kif7-Gli interaction contributes to the organization of the cilium tip compartment and Gli regulation. The lack of mechanistic insights has precluded a deeper understanding of the role of the microtubule cytoskeleton and the primary cilium in Hh signaling. Here we address these fundamental questions and uncover new structural and biochemical design principles by which the cytoskeleton can act as a cytoplasmic scaffold for transcription factor regulation.

## Results

### DNA structural mimicry underlies Gli transcription factor binding to Kif7

Gli proteins were shown to co-immunoprecipitate with Kif7 in pull-down assays from mouse embryo lysates^23^. However, biochemical and structural characterization of Gli-Kif7 complexes has been challenging due to high proteolytic sensitivity and low solubility of these large (>150 kDa) multi-domain proteins. To overcome this technical challenge, we used a pull-down assay to map the interaction domains between Gli and Kif7. We observed that the DNA-binding zinc finger domain of Gli2 (418-594aa; Gli2-ZF) bound to the first coiled-coil dimerization domain of Kif7 (460-600aa; Kif7-CC) **(Fig. 1a,b)**. In addition, a weak signal corresponding to interaction between monomeric Kif7 motor domain (1-361aa) and Gli2-ZF was observed **(Extended Fig 1a)**. We first characterized the interaction between Kif7-CC and Gli2-ZF. Bio-layer interferometry (BLI)-based binding assay with purified recombinant proteins **(Extended Data Fig. 1b,c)** showed tight binding of the proteins in the Kif7-CC:Gli2-ZF complex with a K_d o_f 48 ± 5 nM **(Fig. 1c)**. We were able to further narrow the interaction site on Kif7 to a 63 amino acid shorter coiled-coil segment of Kif7-CC (481-543aa; Kif7-SCC) that binds Gli2-ZF with a similar K_d o_f 69 ± 20 nM **(Fig. 1c)**. Together, these data show that Kif7 uses its coiled-coil dimerization domain, and not the canonical C-terminal ‘cargo’ binding domain in kinesins, to bind Gli2 with high affinity, while Gli2 uses its DNA-binding domain to bind Kif7 dimer. Thus, the domain of the Gli transcription factor that mediates its interaction with DNA in the nucleus is repurposed to interact with a kinesin in the cytoplasm.

**Figure 1.**
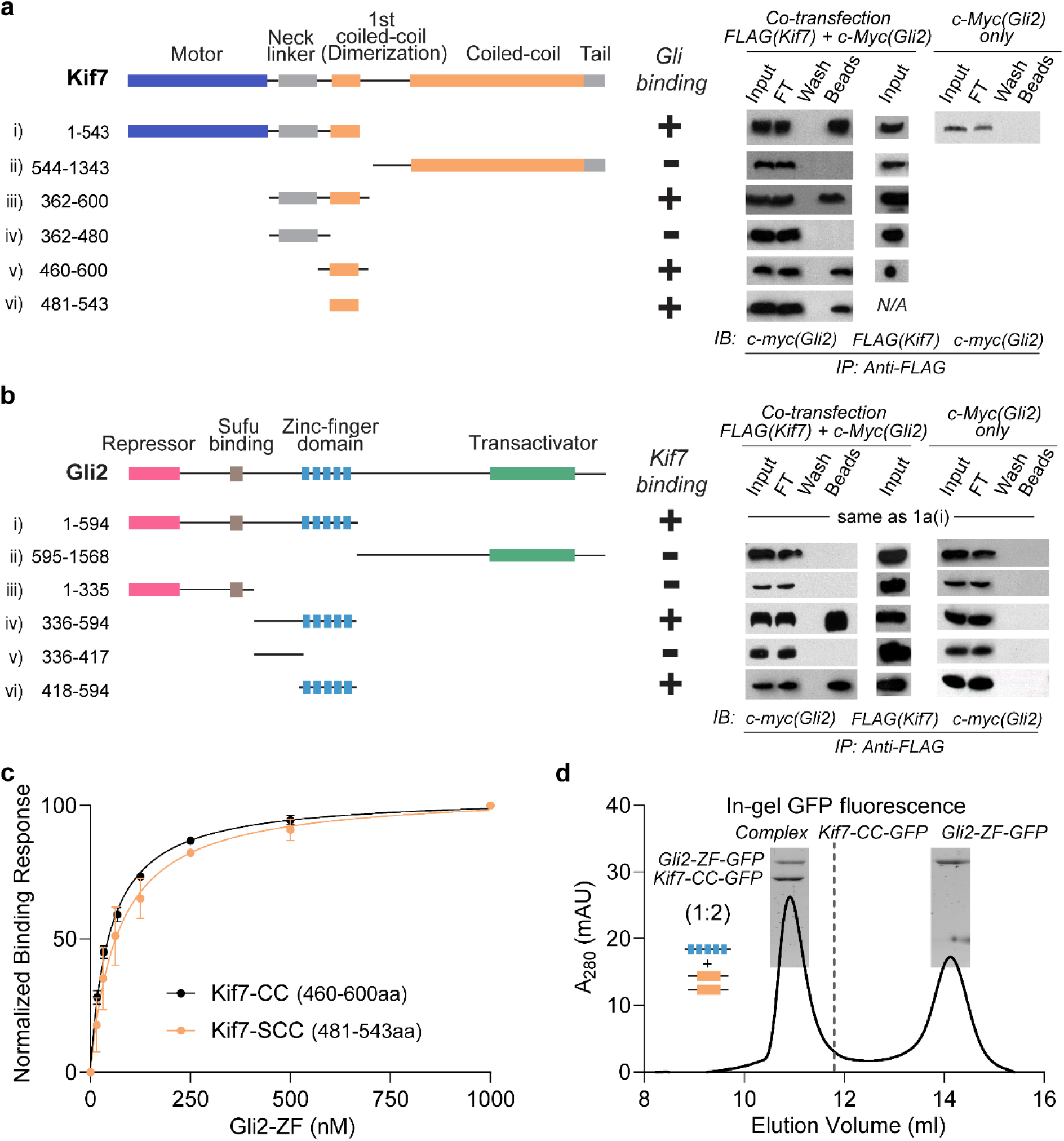
DNA-binding domain of Gli2 forms a high affinity complex with the dimerization domain of Kif7. **a,** Elucidation of the Kif7 domains that interact with Gli2. Pull-down of c-Myc- Gli2(1-594aa) and FLAG-Kif7 constructs (i to vi) after co-transfection in Expi293F cells. *(left panel)* Domain architecture of full length Kif7 and deletion constructs used in this experiment. *(right panel)* Immunoprecipitation (IP) using anti-FLAG magnetic beads. Input (cell lysate), FT (flow through), wash and beads samples were immunoblotted (IB) with anti c-myc antibody to detect Gli2. Transfection with c-Myc-Gli2 (1-594aa) alone was included as a negative control. The blots are representative images of a minimum of three repeats. *(center panel)* Summary of results. The +/- sign represents presence/absence of binding. **b,** Elucidation of the Gli2 domains that interact with Kif7. Pull-down of c-Myc-Gli2 constructs (i to vi) with different FLAG-Kif7 constructs after co-transfection in Expi293F cells. Kif7(1-543aa) was used for i) & ii), Kif7(362- 600aa) was used for iii) & iv) and Kif7(460-600aa) was used for v) & vi). *(left panel)* Domain architecture of full length Gli2 and deletion constructs used in this experiment. *(right panel)* Immunoprecipitation (IP) using anti-FLAG magnetic beads. Input (cell lysate), FT (flow through), wash and beads samples were immunoblotted (IB) with anti c-myc antibody to detect Gli2. Transfection with each of the different c-Myc-Gli2 constructs alone were included as negative controls. The blots are representative images of a minimum of three repeats. *(center panel)* Summary of results. The +/- sign represents presence/absence of binding. **c,** Bio-Layer Interferometry (BLI) assay to quantitatively examine the binding affinity of Kif7 coiled-coil domain constructs; GST-Kif7-CC (460-600aa; black) & GST-Kif7-SCC (481-543aa; orange), to Gli2-ZF. Data represent mean and standard deviation from three independent repeats. The plots of binding response versus Gli2-ZF concentration were fit to a Hill equation to determine equilibrium dissociation constants (K_d)_. For Kif7-CC (460-600aa): K_d =_ 48 ± 5nM, for Kif7-SCC (481-543aa): K_d =_ 69 ± 20nM. **d,** Chromatograms from size exclusion chromatography of a mixture of 125µM Kif7-CC-GFP and 750µM Gli2-ZF-GFP (Superdex 200 10/300 GL). SDS-PAGE of the complex (inset) was used to determine stoichiometry by in-gel GFP fluorescence analysis and is indicated in parenthesis. Gel image is representative of a minimum of three repeats.

How does the DNA binding domain of Gli bind the coiled-coil dimerization domain of Kif7? Single molecule fluorescence intensity and photobleaching analysis of the GFP-tagged proteins indicate that the GST-Kif7-CC is an obligate dimer and Gli2-ZF is a monomer **(Extended Data Fig. 1d,e)**. We determined that the stoichiometry of the Kif7-CC-GFP:Gli2-ZF-GFP complex was 2:1; using in-gel GFP fluorescence and quantitative Western blot **(Fig. 1d and Extended Data Fig. 1f,g;** see Methods). Thus, one molecule of monomeric Gli2-ZF binds to one Kif7-CC dimer.

To gain insight into the structural basis of the Kif7-SCC:Gli2-ZF complex, we performed comparative homology modeling. The structure of Gli2-ZF domain was modelled using the X-ray structure of the zinc-finger domain of Gli1 isoform as a template (PDB: 2GLI, >95% sequence similarity)^34^ **^(^Fig. 2a)**. The Kif7 coiled-coil domain was modeled as a homodimer with high accuracy using X-ray structures of other coiled-coil domains as template **(Fig. 2c)**. The complex between Gli and DNA is characterized by the positively charged zinc-finger domain of Gli wrapping around the major groove of the rod-like DNA double-helix. Extensive electrostatic contacts exist between the negatively charged phosphate backbone of DNA and the second and third zinc-fingers in Gli^34^. As expected, the structural model of Gli2-ZF is very similar to Gli1-ZF **(Extended Fig. 2a-c and Fig 2b)**. Analysis of the structural model of Kif7-SCC revealed two striking features. First, the overall size and shape of Kif7-SCC is similar to dsDNA. Second, analysis of the electrostatic surface potential of Kif7-SCC (pI = 4.48) shows a highly negatively charged rod-shaped surface, similar to the DNA backbone **(Fig. 2c)**. Modelling the structure of the Kif7-Gli2 complex indicates that the positively charged interface of Gli2 zinc-fingers clamp the negatively charged Kif7-SCC rod-like domain **(Extended Data Fig. 2a-c)** in a manner similar to the Gli-DNA complex. The stark similarities in electrostatic surface charge complementarity, molecular dimensions, and mode of binding of Gli2-ZF with DNA and Kif7-SCC suggests that Kif7 may be a DNA molecular mimic for Gli binding in the cytoplasm.

**Figure 2.**
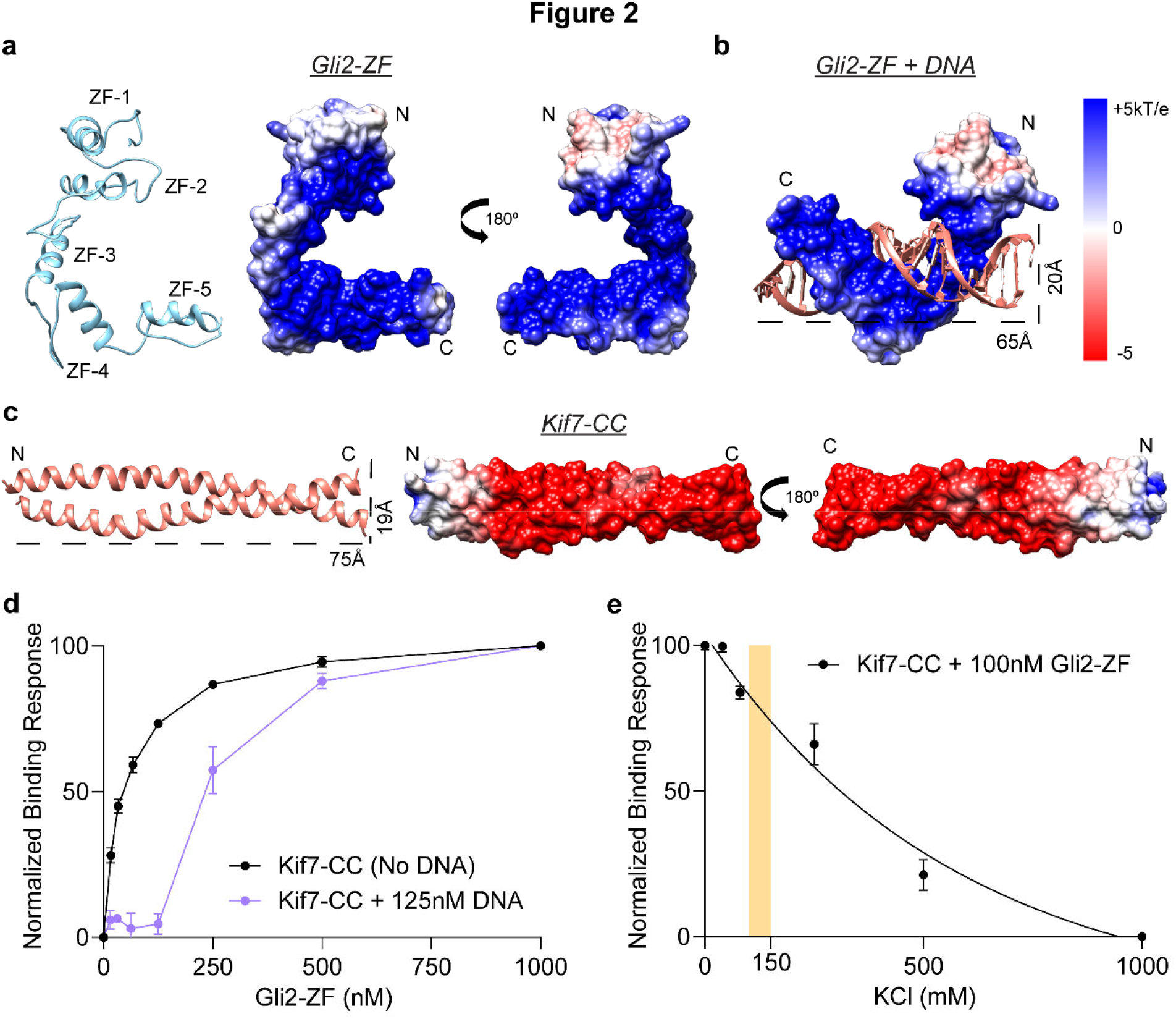
The coiled-coil dimerization domain of Kif7 is a DNA structural mimic for Gli binding. **a,** Structural model of Gli2-ZF (ribbon diagram; SWISS MODEL server: GMQE = 0.98, QMean = -5.14) and overall electrostatic surface representation of the model. **b,** Structural model of Gli2-ZF bound to DNA (based on Gli1-ZF-DNA crystal structure; PDB: 2GLI) included for comparison. **c,** Structural model of Kif7-CC (ribbon diagram; SWISS MODEL server: GMQE = 0.55, QMean = 1.84) and overall electrostatic surface representation of the model. Dotted lines represent molecular dimensions of Kif7-CC and DNA. **d,** Binding of Gli2-ZF and Kif7-CC in the absence (black) and presence of 125nM sequence-specific Gli2 target dsDNA (purple). Normalized binding response from the BLI measurements is plotted against Gli2-ZF concentration. Data represent mean and standard deviation from three independent repeats. **e,** Effect of increasing ionic strength (KCl concentration) on the Kif7-CC and Gli2-ZF binding. Normalized binding response was measured at 100 nM Gli2-ZF using the BLI assay. Data represent mean and standard deviation from three independent repeats. Yellow bar marks physiological ionic strength (∼125-150mM KCl).

To test the structural model of the Kif7-CC:Gli2-ZF interaction we first examined whether Kif7-CC and DNA compete for the same binding site on Gli2. We measured the binding between Gli2-ZF and Kif7-CC in the presence of a Gli-specific target dsDNA (TRE-2S)^35^. In the presence of 125nM TRE-2S DNA there was no detectable binding between Kif7-CC and Gli2-ZF until the analyte concentration was above 125nM Gli2-ZF **(Fig. 2d)**. This suggests that DNA-bound Gli2- ZF cannot bind Kif7-CC and that these interactions are mutually exclusive. We conducted the BLI binding assay in buffers with increasing ionic strength and observed a systematic reduction in the measured binding response from Kif7-CC and Gli2-ZF interaction, confirming that the interaction between these proteins is driven by electrostatic interactions **(Fig. 2e)**.

Finally, we performed a series of site-directed mutagenesis experiments to further confirm the structural model and map the residues of Gli2-ZF and Kif7-CC required for complex formation. BLI binding assay with Gli2-ZF truncations containing different groups of zinc-fingers indicated the necessary requirement for zinc-fingers 2 and 3 in tandem for Kif7-CC binding **(Extended Data Fig. 3a-d)**. This is similar to the Gli-DNA interaction where ZF2 and ZF3 are critical for making contacts with the phosphate backbone of DNA^34^. Mutation of specific charged residues in Gli2 ZF-2 (H493A/R496A) and ZF-3 (H503A/R516A/K521A) abolished binding to Kif7-CC whereas those in ZF-4 (R550A/K552A/R556A) did not affect binding **(Extended Data Fig. 3e,f)**. Interestingly, the residues H493 (ZF2), R516 (ZF3) and K512 (ZF3) also engage in phosphate backbone contacts with DNA^34^, suggesting that the same subset of amino acids interact with Kif7- CC. Analysis of the interaction surface on Kif7-CC revealed three patches of negatively charged amino acid residues (labeled as S1, S2 and S3 in **Extended Data Fig. 3f**) that are potential Gli2- ZF binding residues. Systematic mutation of residues in these patches revealed that while Kif7- CC S1 (E500A/E501A/E502A/D505A) mutant abolished binding to Gli2-ZF, the S2 (E511A/E515A) and S3 (E526A/E529A & E530A/R535A) mutants did not (**Extended Data Fig. 3e,f)**. To further validate the interaction, we expressed and purified four recombinant S1 mutants (D505A, E501A/E502A, E502A, E502A/D505A). All four mutants elute similar to wild type protein in size exclusion chromatography, indicating that they are properly folded **(Extended Data Fig. 3g,h)**. Three of these mutants (E501A/E502A, E502A, E502A/D505A) showed no binding to Gli2-ZF while the fourth one (D505A), shows diminished binding **(Extended Data Fig. 3i)**. This suggests that negatively charged residues included in S1 that lie approximately at the center of the coiled-coil rod of Kif7 form the primary contact point for Gli binding.

Taken together, the results from DNA competition assay, ionic strength-dependence and the mutagenesis studies provide additional support for DNA mimicry by the coiled-coil domain of Kif7. The center of this negatively-charged rod forms the Gli2 interaction site with zinc-fingers 2 and 3 making key charge-charge contacts, as seen previously in binding of Gli to DNA^34^.

We characterized the interaction between Gli2-ZF and the dimeric Kif7. We determined that the stoichiometry of Kif7-DM:Gli2-ZF complex in the absence of microtubules was 1:1, suggesting there are two molecules of Gli2-ZF bound per Kif7-DM **(Fig. 3a and Extended Data Fig. 4a-e)**. Since one Gli2-ZF is bound to the coiled-coil dimer of Kif7, we hypothesized that the second Gli2-ZF molecule binds proximal to the Kif7 motor domains, as was also suggested by the weak signal observed in the pull-down assay using cell lysate (**Extended Data Fig. 1a)**. However, we failed to detect any binding between the purified recombinant monomeric Kif7 motor domain (1-386aa, Kif7-MM) and Gli2-ZF at concentrations as high as 12µM Gli2-ZF in the BLI assay. The differences in the two experiments could arise from the presence of low levels of intact microtubules in the cell extracts. We therefore examined the recruitment of Gli to GMPCPP-polymerized microtubules by Kif7 using an *in vitro* TIRF microscopy assay (**Fig. 3b)**. We examined the microtubule localization of Alexa-647 labeled Gli2-ZF-SNAP in the presence of a monomeric Kif7 motor domain construct (1-386aa, Kif7-MM-GFP), which lacks the coiled-coil Gli interaction site, and compared it to dimeric Kif7 protein (1-543aa, Kif7-DM-GFP), which has both sites. Analysis of the Alexa-647 intensity reveals microtubule-bound Kif7-MM can bind Gli2- ZF, albeit to a lesser extent when compared to the Kif7-DM (∼9 fold versus ∼3 fold above the background fluorescence of Gli alone on microtubules respectively) **(Fig. 3c)**. These data suggest that the motor domain of Kif7 likely forms a Gli-interaction site, but this interaction requires concentrating motors on microtubules or another mechanism, such as dimerization, that brings the motor domains in proximity.

**Figure 3.**
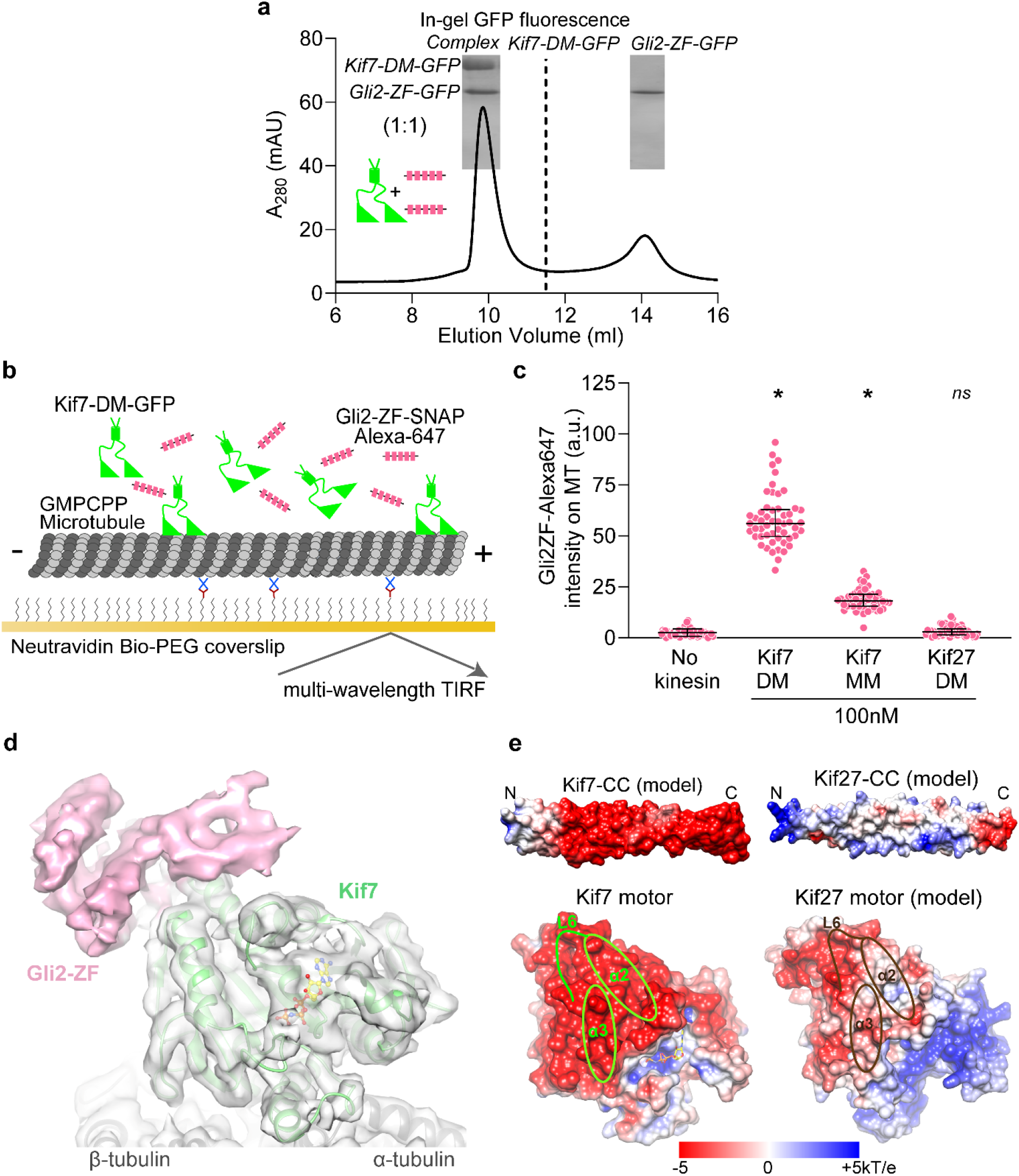
Second Gli interaction site on Kif7 motor domain. **a,** Chromatograms from size exclusion chromatography of a mixture of 125µM Kif7-DM-GFP and 750µM Gli2-ZF-GFP (Superdex 200 10/300 GL). SDS-PAGE of the complex (inset) was used to determine stoichiometry by in-gel GFP fluorescence analysis and is indicated in parenthesis. Gel image is representative of a minimum of three repeats. **b,** Schematic of the *in vitro* total internal reflection fluorescence (TIRF) microscopy-based assay used to examine microtubule-localized Gli2-ZF and Kif7-DM. Rhodamine/HiLyte 647 labeled GMPCPP-stabilized microtubules (grey) were immobilized on a PEG-treated glass coverslip (yellow) via neutravidin-biotin linkages (blue & brown respectively). Kif7-DM-GFP (green), Gli2-ZF-SNAP-Alexa 647 (pink) and 1mM ATP were subsequently added to examine binding of both proteins on microtubules. **c,** Scatter plot of Gli2-ZF-Alexa647 intensity per pixel on microtubules (MT) represents recruitment of Gli on MT by Kif7-MM and Kif7-DM. The no kinesin condition was a control for non-specific binding of Gli2-ZF to MT and Kif27-DM was included as a negative control for Gli2-ZF binding. Assay conditions: 100nM Gli2-ZF-SNAP-Alexa647 with 100nM of kinesin in each case. N ≥ 50 microtubules for each condition. One-way ANOVA (*p* < 0.0001) and post-hoc analysis (\**p* < 0.0001 in Dunnett’s multiple comparisons test) show statistically significant differences in Gli intensity compared to no kinesin control; *ns* is not significant (*p* = 0.9253). **d,** Cryo-EM reconstruction of Kif7-DM bound to microtubules in the presence of Gli2-ZF-SNAP and AMPPNP. The structural model for Kif7 motor domain (green ribbon) bound to microtubule (grey ribbon) was fitted in the density and refined. The density corresponding to Gli2-SNAP is shown in pink. AMPPNP is shown as a yellow ball-and-stick model. **e,** Comparison of electrostatic surface potential of coiled-coil and motor domains of Kif7 (PDB: 6 MLR) and Kif27. The structural models of Kif7-CC (SWISS MODEL server: GMQE = 0.55, QMean = 1.84), Kif27-CC (SWISS MODEL server: GMQE = 0.51, QMean = 0.84) and Kif27 motor domain (SWISS MODEL server: GMQE = 0.47, QMean = -2.22) were obtained by homology modelling. Loop L6, helices α-2 and α-3 on the Kif7 motor domain that lie close to the Gli2-SNAP density are highlighted in green. Corresponding regions (Loop L6, helices α-2 and α-3) in the Kif27 motor domain model are highlighted in brown.

To visualize the binding of Gli2-ZF to the Kif7 motor domain we obtained cryo-EM reconstructions of Kif7-DM bound to microtubules in the presence of Gli2-ZF-SNAP and the non-hydrolysable ATP analogue, AMP-PNP (3.9 Å resolution, **Fig. 3d, Extended Data Fig. 5a,b and Extended Data Table 1)**. Comparisons of the cryo-EM reconstruction with a map of microtubule-bound Kif7 motor domain in the AMP-PNP state (PDB:6MLR, EMD:9141) showed that the motor domain adopts the same conformation while binding to microtubules in the presence and absence of Gli2-ZF **(green in Fig. 3d)**. While no additional density was seen bound to the microtubule lattice, an extra region of density contacting the motor domain of Kif7 was observed in the presence of Gli2-ZF-SNAP **(pink in Fig. 3d)**. Close inspection reveals that this extra density lies close to loop L6, helices α-2 and α-3 on the Kif7 motor domain and away from the nucleotide-binding pocket as well as the microtubule-binding interface of the motor **(Supplementary Video 1)**. We confirmed that the extra density originated from the Gli2-ZF protein by generating and comparing cryo-EM reconstructions of the following microtubule-bound complexes: (i) Kif7-MM in the AMP-PNP state with Gli2-ZF-SNAP at 4.8 Å resolution, and (ii) Kif7-DM in the ADP state with Gli2-ZF, at 4.3 Å resolution **(Extended Data Table 1).** The presence of extra density in the Kif7-MM:Gli2-SNAP map at the same position as in Kif7-DM:Gli2-SNAP map ruled out the contribution of Kif7-coiled-coil and neck-linker domains to the density **(Extended Data Fig. 5c)**. Comparison of the Kif7-DM:Gli2-ZF and Kif7-DM:Gli2-SNAP maps confirmed that a large part of the extra density arose from Gli2-ZF and not SNAP **(Extended Data Fig. 5d)**. Superposition of the cryo-EM maps obtained in the presence of Gli2-ZF and Gli2-ZF-SNAP was used to unambiguously assign the density corresponding to Gli2-ZF **(Extended Data Fig. 5d)**. We built a structural model of the Gli2-ZF peptide backbone sequence (see Methods) into this extra density and were able to fit two complete and one partial zinc-fingers **(Extended Data Fig. 5e and Supplementary Video 1)**. Since it is well established that under saturating motor concentrations needed for cryo-EM reconstructions, the density of only a single motor head can be resolved^36, 37^, we do not expect to see the entire Gli2-ZF interaction site formed by both motor heads and the coiled-coil domain in a dimer. TIRFM assays with Alexa-647 tagged Gli2-ZF1-3-SNAP protein show that first three ZFs of Gli are sufficient to bind Kif7-MM **(Extended Data Fig. 4f)**. Together these results confirm that in addition to the high affinity interaction site on Kif7-CC, there is a second interaction site for Gli2-ZF proximal to the Kif7 motor domain. Kif27 is a close homolog of Kif7 and these two proteins are thought to arise from a gene duplication event^38, 39^. Unlike Kif7, Kif27 plays a role in motile cilium biogenesis and is not thought to be involved in Hedgehog signaling^40^. We wondered if the structural determinants of Gli interaction sites are conserved or dissimilar between Kif7 and Kif27. We modelled the motor domain (1-398aa) and the coiled-coil dimerization domains (480-550aa) of Kif27 and Kif7 and compared their electrostatic surface potential. Our analysis revealed that the electrostatic surface potential of Kif27-CC is overall neutral in contrast to Kif7-CC, and the motor domain regions of Kif27 (helix α-2, α-3 and loop L6) corresponding to the Gli2-ZF interaction site in Kif7, present a less negative surface potential **(Fig. 3e)**. Consistent with this, we do not observe recruitment of Gli2-ZF on microtubules by the dimeric Kif27-DM-GFP (1-580aa) protein in our TIRF-microscopy reconstitution assay **(Fig. 3c)**. The comparative structural analysis of Kif7 and Kif27 indicates a specific adaptation in Kif7 for Gli binding and Hedgehog signaling.

### Synergistic localization of Kif7 and Gli to microtubules *in vitro*

We next examined if the binding of Gli altered the microtubule-binding of Kif7. The microtubule binding of Kif7-DM-GFP was imaged in the absence or presence of Alexa-647 labeled Gli2-100nM ZF-SNAP in a TIRFM assay. As expected, analysis of Alexa-647 intensity showed 3-fold increase in recruitment on microtubules in the presence of Kif7-DM **(Fig. 4a)**. Low levels of Gli2 fluorescence intensity were visualized on microtubules even in the absence of Kif7, which occurs from non-specific electrostatic interactions in lower ionic strength BRB80 buffer. Interestingly, analysis of Kif7 fluorescence intensity (100nM) revealed a >5-fold increase in Kif7 binding to microtubules in the presence of Gli2-ZF (50nM) **(Fig. 4b)**. Thus, Gli binding to Kif7 changes the microtubule binding affinity of Kif7 and sets up a positive feedback loop that in turn results in enhanced recruitment of Gli to microtubules. This positive feedback in the kinesin-microtubule binding generates a graded response that is sensitive to increasing Gli concentrations until it reaches saturation **(Fig. 4c)**. Further examination of the Kif7-microtubule interaction revealed an increase in the residence time of single molecules of Kif7-DM-GFP on microtubules with increasing concentrations of Gli2-ZF. **(Extended Data Fig. 6a-c)**. We validated the findings from TIRF assays using a co-sedimentation assay, which showed that Kif7 has a higher microtubule binding affinity in the presence of Gli2-ZF (K_d =_ 0.54 ± 0.03 μM) compared to when Gli is absent (K_d =_ 4.23 ± 0.24 μM) **(Fig. 4d and Extended Data Fig. 6d)**.

**Figure 4.**
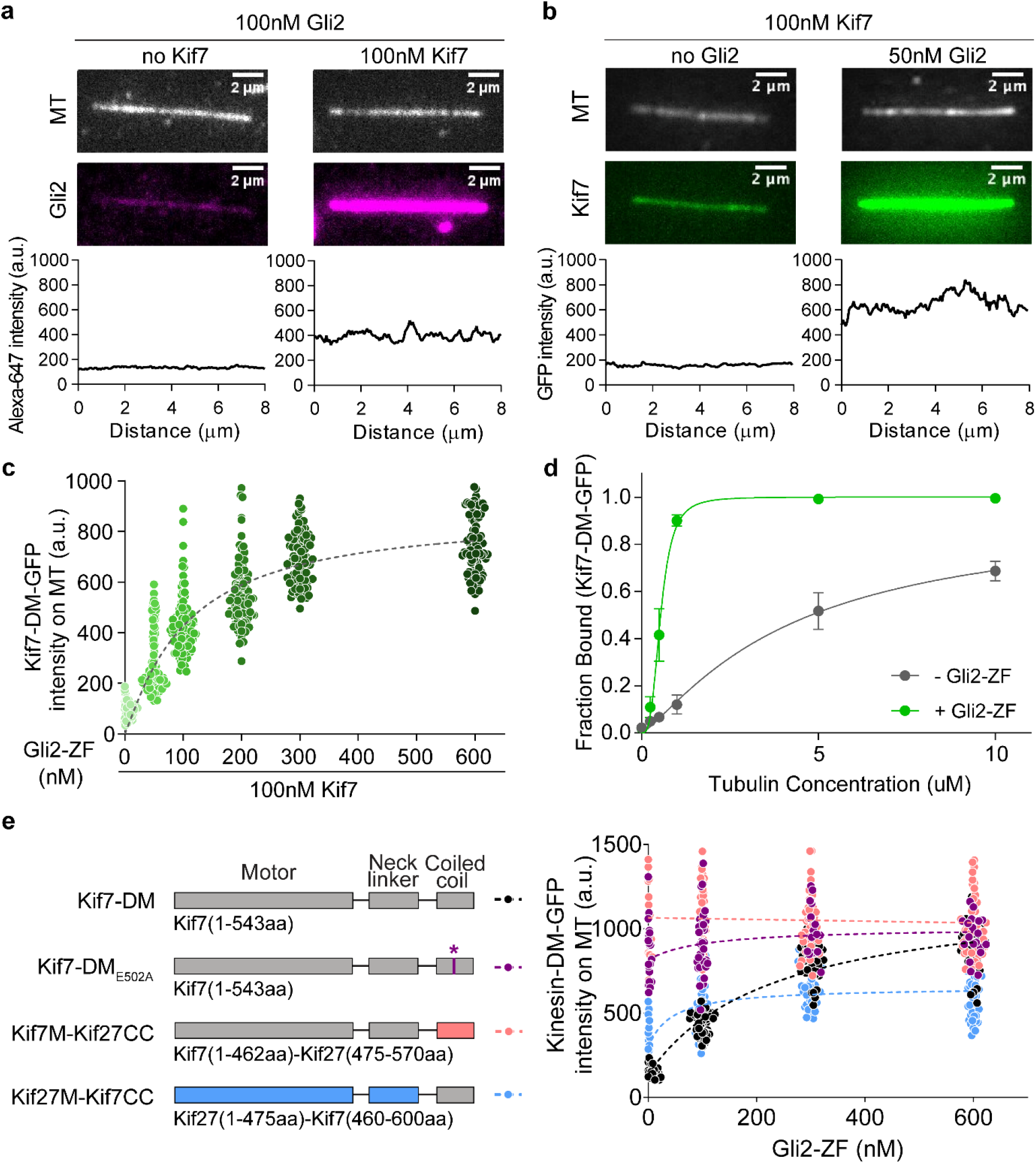
Gli2 ZF increases the microtubule-binding affinity of Kif7, thereby increasing its own recruitment on microtubules. **a,** Representative images of microtubule (MT, top) and Gli2- ZF-Alexa 647(Gli2, bottom, 100nM) in the absence (left) or presence of Kif7-DM-GFP (right, 100nM). The graphs below show line scans of the corresponding Alexa 647 fluorescence intensity. Scale bars represent 2μm. **b,** Representative images of microtubule (MT, top) and Kif7-DM-GFP (Kif7, bottom, 100nM) in the absence (left) or presence of Gli2-ZF (right, 50nM). The graphs below show line scans of the corresponding GFP fluorescence. Scale bars represent 2μm. **c,** Scatter plot of Kif7-DM-GFP intensity per pixel on microtubules (MT) in the presence of increasing concentrations of Gli2-ZF. Assay conditions: 100nM Kif7-DM-GFP with 0, 50, 100, 200, 300 & 600nM Gli2-ZF. N ≥ 90 microtubules at every Gli2-ZF concentration. One-way ANOVA (*p* < 0.0001) and post-hoc analysis (*p* < 0.0001 in Dunnett’s multiple comparisons test) shows statistically significant differences among means at various Gli2-ZF concentrations compared to no Gli2-ZF. Data fit to dose response curve (dotted grey line) with half maximal response at ∼160 nM Gli2-ZF. **d,** Co-sedimentation assay curve from analysis of gels in **c,** to quantitatively examine the microtubule binding affinity of Kif7DM-GFP in the absence (grey) and presence (blue) of Gli2-ZF. Error bars represent standard error. The plots of fraction of Kif7DM-GFP bound versus microtubule concentration were fit to a Hill equation to determine the equilibrium dissociation constants (K_d)_. For Kif7-DM-GFP - Gli2-ZF: K_d =_ 4.23 ± 0.24 μM (R^2=^ 0.93) and for Kif7-DM- GFP + Gli2-ZF: K_d =_ 0.54 ± 0.03 μM (R^2=^ 0.95). In the presence of Gli2-ZF, Kif7-DM-GFP shows co-operative binding to microtubules with a Hill co-efficient (h) of 3.2 ± 0.59. **e,** Schematics showing the domains swapped in the Kif7-Kif27 chimera proteins and the E502A point mutant Kif7-DM. Scatter plot of Kinesin-GFP intensity per pixel on microtubules (MT) in the presence of increasing concentrations of Gli2-ZF. Assay conditions: 100nM of each kinesin: Kif7-DM (black), Kif7-DM_E502A (_purple), Kif7M-Kif27CC (red) and Kif27M-Kif7CC (blue), with 0, 100, 300 & 600nM Gli2-ZF. N ≥ 40 microtubules for each data set. Two-way ANOVA (factor 1: Gli2-ZF concentration *p* < 0.0001; factor 2: kinesin type *p* < 0.0001 and their interaction *p* < 0.0001) and post-hoc analysis (*p* < 0.01 in Dunnett’s multiple comparisons test) shows statistically significant differences among means at various Gli2-ZF concentrations compared to Kif7-DM. Data were fit to dose response curve (dotted lines).

We wanted to determine the contribution of the two Gli interaction sites on Kif7 in regulating the Kif7-microtubule interaction. We first hypothesized that the binding of Gli2-ZF to the Kif7 motor domain may underlie the observed changes in the Kif7-microtubule binding. Therefore, we generated a chimera in which the Kif7 motor domains are intact, but the Kif7-CC is replaced by Kif27-CC (Kif7M-Kif27CC) **(Fig. 4e and Extended Data Fig. 7a,b)**. To our surprise the Kif7M-Kif27CC chimera has constitutive higher binding to microtubules compared to Kif7-DM and the kinesin-microtubule interaction was independent of Gli concentration **(Fig. 4e)**. To test this further we generated a point mutation in Kif7-DM (E502A), which eliminates Gli binding to the coiled-coil domain of Kif7 **(Extended Data Fig. 3e)**. Similar to Kif7M-Kif27CC, Kif7-DM(E502A) is a constitutive dimer and recruits lower levels of Gli2-ZF relative to wild type as expected for constructs lacking one Gli binding site **(Extended Data Fig. 7)**. We find that this mutant is identical to the Kif7M-Kif27CC chimera in terms of Gli-independent strong microtubule binding **(Fig.4e)**. Thus, counterintuitive to our expectation, the binding of Gli2-ZF to the Kif7-CC domain rather than the Kif7 motor domain has a dominant role in the regulation of Kif7- microtubule interaction. These data are consistent with a mechanism where the coiled-coil domain inhibits the Kif7-microtubule binding, and this inhibition is relieved by Gli binding to the coiled-coil and motor domains.

**Figure 7.**
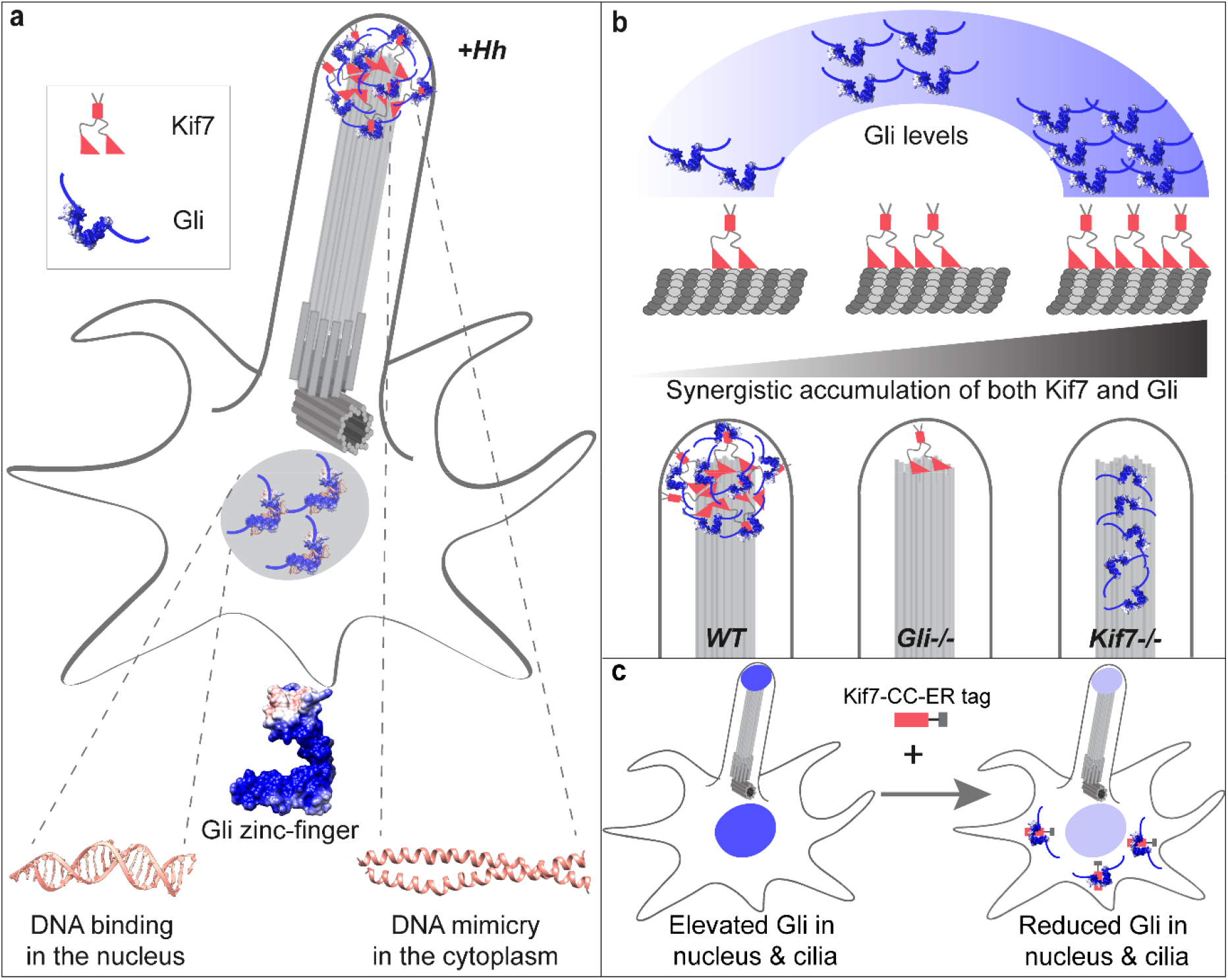
Schematic of the structural basis and cellular implications of Kif7-Gli interaction. **a,** DNA molecular mimicry by the Kif7 coiled-coil underlies the tethering of Gli in the cytoplasm and the cilia tips. The zinc-finger domain of Gli that binds DNA in the nucleus is co-opted for binding the Kif7 coiled-coil out of the nucleus. **b,** The Kif7 coiled-coil acts as a regulatory domain necessary for the graded accumulation of both proteins on microtubules and at the cilium tip in a Gli-concentration responsive manner. Consistent with this, Gli knockout cells (Gli-/-) have low levels of Kif7 at the cilia tips compared to WT cells during Hedgehog signal transduction. In Kif7 knockout cells (Kif7-/-) Gli is distributed in puncta along the length of the ciliary axoneme and is not concentrated at the cilia tip during pathway activation^25, 30^. **c,** The Kif7 coiled-coil peptide can be re-tooled as a reagent for sequestration of Gli away in the cytoplasm to decrease nuclear and ciliary Gli levels.

To test if the inhibition by the coiled-coil is specific to the Kif7 motor domain, we generated a chimera where the motor and neck linker domain of Kif7 was replaced by that of Kif27 (Kif27M-Kif7CC). Similar to the other mutants described in this section, this chimera is dimeric and recruits lower levels of Gli compared to wild type Kif7-DM **(Extended Data Fig. 7)**. We found that the microtubule binding of Kif27M-Kif7CC chimera exhibited lower dependence on Gli concentrations **(Fig. 4e and Extended Data Fig. 7)**. Together these data suggest that the inhibition of the Kif7-microtubule binding by the Kif7 coiled-coil domain is highest in the context of the Kif7 motor domains in the Kif7-DM, and the complete alleviation of this inhibition by Gli sets the dynamic range of the Kif7-microtubule binding in response to changes in Gli concentration.

### Synergistic localization of Kif7 and Gli2 at the distal cilia tips

Our pull-down and in vitro experiments suggested that Kif7 can interact with both the N-terminal repressor as well as the zinc finger domain of Gli2. We examined the interaction between full-length Gli2 and Kif7 in the cellular context. When overexpressed individually Kif7-FL is exclusively localized in the cytoplasm and Gli2-FL is largely in the nucleus with a small fraction localized in cytoplasmic puncta and not on microtubules. Co-transfection of Kif7-FL-mRuby with mNeonGreen-Gli2-FL in HeLa cells revealed localization of both Kif7 and Gli2 along filamentous microtubules in the cytosol **(Fig. 5a)**. Kif7 and Gli2 are known to accumulate at the cilium tip in response to Hedgehog (Hh) pathway activation, and Kif7 is thought to form a stable platform essential for proper recruitment of Gli to the cilium tip^25, 30^. Contrary to this model, our biochemical analysis of the Kif7-Gli interaction on microtubules suggest that the cilium localization of Kif7 may in turn be Gli-dependent. To examine this, we measured the amount of Kif7 at the cilium tip in Gli2-/- and Gli2-/-Gli3-/- cell lines (immortalized MEFs) compared to wild-type (WT) MEFs, upon stimulation with the Smoothened agonist SAG (Hh pathway agonist). We found that the amount of Kif7 at cilia tips during Hh pathway activation depends on Gli. Gli2-/- cells showed ∼4-fold less Kif7 in the cilia, whereas Gli2-/-Gli3-/- cells showed 5-fold less Kif7 at the cilia tips compared to WT cells **(Fig. 5b,c)**. Thus, by modulating the level of Kif7 at the cilium tip, Gli acts as a master regulator of its own recruitment to the cilia tip for Hh pathway activation. Together these findings suggest that the Kif7-Gli2 interaction is not only sufficient to recruit Gli onto microtubules, but it also positively regulates the microtubule-binding of Kif7 by increasing the amount of motor on microtubules and at the cilium tip. This positive regulation in turn increases the amount of Gli2 recruited to microtubules. Consequently, the interaction between Gli2 and Kif7 concentrates both proteins to higher levels on microtubules in a Gli-concentration dependent manner.

**Figure 5.**
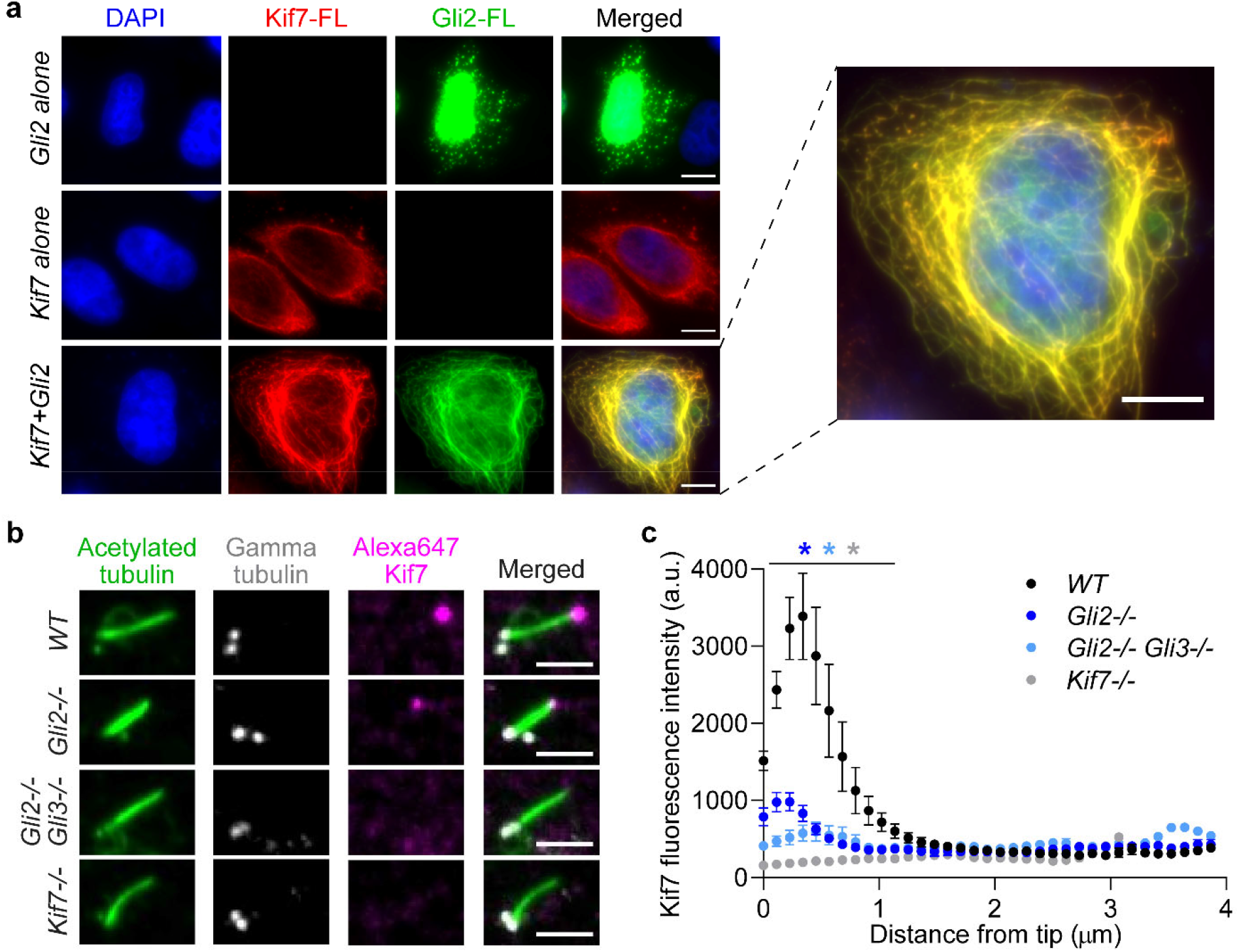
Synergistic regulation of Gli-Kif7 promotes Kif7 localization to cytosolic microtubules and the distal cilium tip. **a,** Localization of transfected mRuby-tagged full length Kif7 and mNeonGreen-tagged full length Gli2 in HeLa cells. Representative images are shown. DAPI staining was used to mark the nucleus. Overexpressed full length Gli2 shows predominant localization in the nucleus (row 1). Overexpressed full length Kif7 is cytoplasmic (row 2). Co- transfection with Kif7 and Gli2 shows localization of both proteins on microtubules in the cytoplasm (row 3) (100%; n>30). Scale bars represent 10 µm. **b,** Cilium-tip localization of Kif7 in *WT, Gli2-/-, Gli2-/-Gli3-/- and Kif7-/-* MEFs. Representative immunofluorescent images of primary cilia are shown. Acetylated α-tubulin antibody was used to mark cilia, γ-tubulin antibody was used to mark centrioles (base of cilia) and Alexa-Fluor-647-labeled Kif7 antibody was used to measure Kif7 amounts in the cilia tips during Hh activation (+SAG). Scale bar represents 2 µm. **c,** Quantitative analysis of the levels of Kif7 in *WT, Gli2-/-, Gli2-/-Gli3-/- and Kif7-/-* MEFs. Line scans of cilia from the experiment in (b), was used to quantify distribution of Kif7 along the length of the cilia. Data represent mean and standard error from three independent repeats (N = 10 cilia for each genotype). Two-way ANOVA (factor 1: genotype *p* < 0.0001, factor 2: distance from tip *p* < 0.0001) and post-hoc analysis (\**p* < 0.0001 in Dunnett’s multiple comparison test) show statistically significant differences among Kif7 intensity in cilia tip of all genotypes compared to wild type control.

### Re-engineering the DNA-mimicking coiled-coil domain of Kif7 as a tool for sequestration of**Gli away from the nucleus**

Small coiled-coil domains and helical peptides have the potential to be developed as inhibitors or cellular sequestration tools to tune the levels of a transcription factor in the nucleus. Given the high affinity of the DNA mimicking Kif7-CC domain for Gli2, we wondered if this domain could be retooled to sequester Gli away from the nucleus^41–43^. To test this possibility, we engineered the Kif7-CC to contain an endoplasmic reticulum (ER) retention tag and mRuby at the C-terminus and expressed it along with full-length mNeonGreen-Gli2 from a single mRNA in a dual expression vector. The identical vector expressing mNeonGreen-Gli2 alone served as a control. These constructs were transfected in HeLa cells and the amount of Gli2 in the nucleus was measured as a percentage of total Gli expression levels using the mNeonGreen fluorescence intensity. The localization of mNeonGreen-Gli2 in control experiments was largely nuclear (>75%) whereas, in the presence of ER-localized Kif7-CC-mRuby the nuclear mNeonGreen-Gli2 decreased to <25 % **(Fig. 6a,b)**. Colocalization of fluorescent signals for Gli2 and Kif7-CC showed that ER-linked Kif7-CC-mRuby sequesters Gli2 in the cytoplasm. We also found that the sequestration of Gli2 in the cytoplasm (and thus away from the nucleus) depends on the level of Kif7-CC-mRuby expression in these cells. Cells with higher Kif7-CC expression showed lower levels of nuclear Gli2 and hence higher levels of cytoplasmic Gli2 sequestration **(Fig. 6c)**. To test if the ER-tagged Kif7-CC could inhibit the cilium tip localization of Gli2 in response to Hedgehog pathway activation we transfected this construct into NIH3T3 cells. The cilium tip localization of Gli2 in Kif7-CC-ER transfected cells, induced with the Hedgehog pathway agonist SAG, was measured using a directly labeled Gli2 antibody that was further cleaned using Gli2-/- MEFs. Our results show that the Kif7-CC-ER inhibits the levels of Gli2 at the cilia tips during pathway activation compared to non-transfected wild type cells **(Fig. 6d,e)**. These findings show that the Kif7-CC peptide can be re-engineered for sequestration of endogenous and over-expressed Gli in the cytoplasm and inhibit its nuclear and cilium localization.

**Figure 6.**
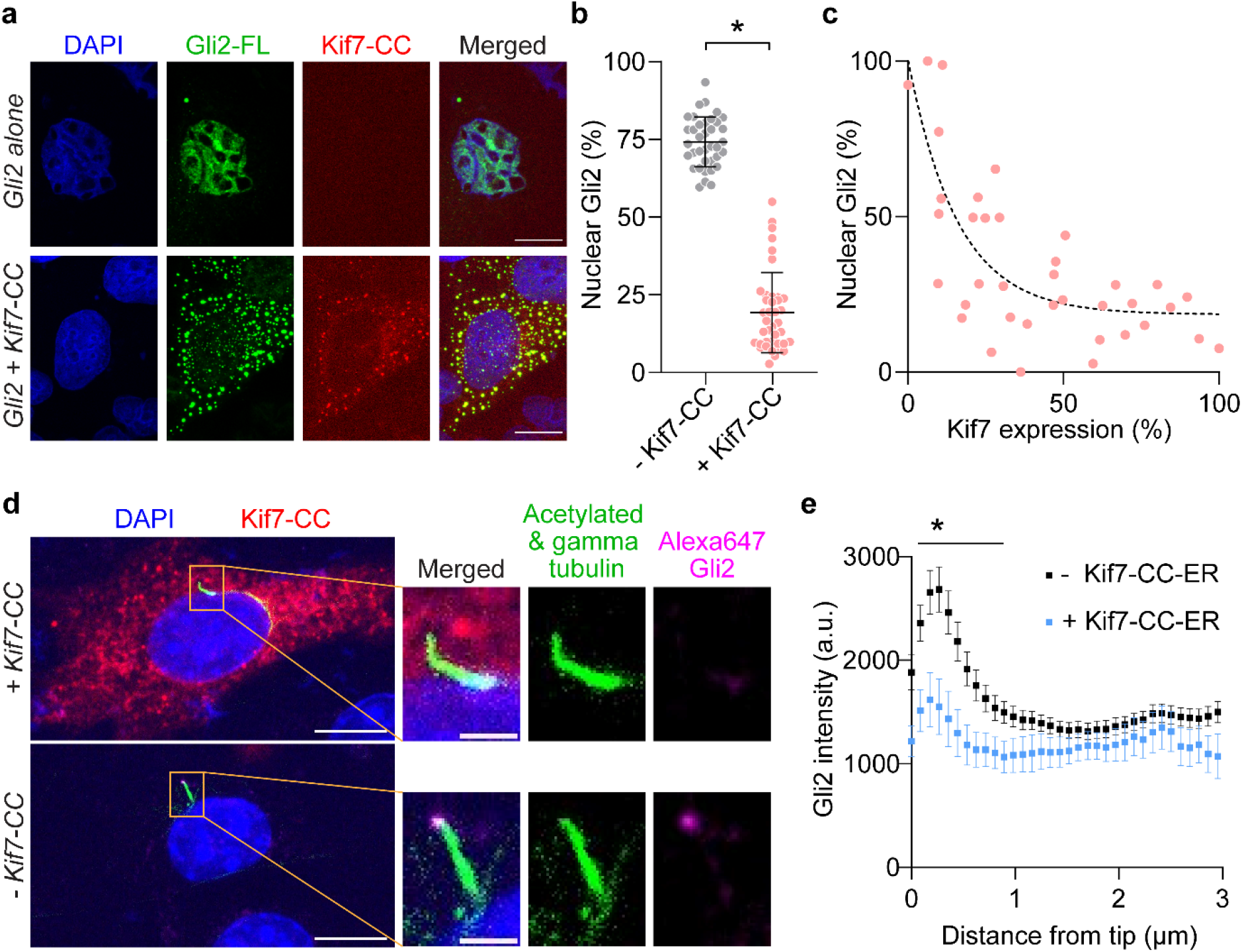
Re-engineering the DNA-mimicking coiled-coil domain of Kif7 as a tool for sequestration of Gli away from the nucleus and cilia. **a,** Localization of transfected mRuby-tagged Kif7-CC (460-600aa) with ER retention tag and mNeonGreen-tagged full length Gli2 in HeLa cells. Representative images are shown. DAPI staining was used to mark the nucleus. Overexpressed full length Gli2 shows predominant localization in the nucleus (row 1). Co-transfection with Kif7-CC and Gli2 shows co-localization of Gli2 in the cytoplasm with Kif7-CC (row 2). Scale bars represent 10 µm. **b,** Percent of nucleus-localized Gli2 as quantified from **a,** in the absence and presence of Kif7-CC. Data represent mean and standard deviation from three independent repeats (N ≥ 35 for each condition). **c,** Scatter plot of percent nuclear Gli2 versus level of Kif7-CC expression. The dotted line represents fit of the data to one-phase association. **d,** Representative immunofluorescent images of cilium-tip localization of endogenous Gli2 in NIH3T3 cell transfected with mRuby-tagged Kif7-CC (460-600aa) with ER retention tag (top row) and untransfected cell from the same sample (bottom row). Acetylated α-tubulin antibody was used to mark cilia, γ-tubulin antibody was used to mark centrioles (base of cilia) and Alexa-Fluor- 647-labeled Gli2 antibody was used to measure endogenous Gli2 amounts in the cilia tips during Hh activation (+SAG). Scale bar in the left panel represents 10µm and in the cilium inset represents 2 µm. **e,** Quantitative analysis of the levels of Gli2 in cilia of NIH3T3 cells transfected with Kif7- CC-ER in the presence of SAG. Line scans of cilia from **d,** was used to quantify distribution of Gli2 along the length of the cilia. Data represent mean and standard error from three independent repeats (N ≥ 12 cilia for each group). Two-way ANOVA (factor 1: transfected with Kif7-CC-ER *p* < 0.05, factor 2: distance from tip *p* < 0.0001) and post-hoc analysis (* *p* < 0.0001 in unpaired Student’s t-test) shows statistically significant differences in Gli2 intensity in cilia tips upon Kif7- CC-ER transfection compared to non-transfected control.

Our study reports the discovery of DNA-mimicry as a mechanism for regulating nuclear proteins in the cytoplasm of eukaryotes and provides critical insights into mechanisms that link the Hh signaling pathway and the microtubule cytoskeleton.

Coiled-coils are among the most ubiquitous structural folds in proteins^44^. The Kif7-CC is characterized by a highly negatively charged rod-like structure that competes for the same binding sites on Gli2-ZF as DNA. Thus, the properties of Kif7-CC are consistent with that of a DNA-mimicking protein^45–48^. The zinc-finger domain of Gli2 binds DNA in the nucleus and is repurposed for Kif7-mediated cytoplasmic localization through the same structural design principle **(Fig. 7a)**. DNA mimicry by the Kif7 coiled-coil is the first example of a coiled-coil domain in an endogenous eukaryotic protein used to bind the DNA-binding domain of a transcription factor and regulate its function in the cytoplasm. A handful of examples of DNA mimicking proteins described in eukaryotes are confined to the nucleus^49, 50^ °r function during mitosis ^51^. Additionally, unlike reported DNA mimics that resemble bent/distorted/single stranded DNA, Kif7-CC mimics dsDNA^49, 50, 52–54^. Our observations raise the intriguing possibility that cytoskeletal proteins can be integral in regulating other nuclear proteins, especially transcription factors, in the cytoplasm, by mimicking double stranded DNA.

We find that Gli binds to two unusual cargo-binding sites on Kif7 - the coiled coil dimerization domain and the motor domain, and the Kif7-Gli interaction regulates the Kif7- microtubule affinity in a Gli dose-dependent manner **(Fig. 7b)**. Experiments with chimeras and mutant that specifically perturb Gli binding to either the coiled-coil or the motor domain in a Kif7- DM provide mechanistic insights into how Gli may regulate the Kif7-microtubule interaction. Our results suggest that in the absence of Gli, the coiled-coil domain of Kif7 may inhibit the Kif7- microtubule interaction. Gli binding to the coiled-coil interaction site on Kif7 alleviates this inhibition. The maximum dynamic range of the Kif7-microtubule binding response to Gli is only observed in the context of the wild type Kif7-DM as replacing the motor domain of Kif7 with that of Kif27 reduces the response. Thus, of the two sites on Kif7 that play a role in recruiting Gli, it is unexpectedly the Gli binding site further away from the motor domain that has a dominant role in Kif7-microtubule binding response. In general, cargo binding to the coiled-coil or the motor domain of kinesins is rare. Particularly, cargo-motor domain interactions typically have an inhibitory effect on the kinesin-microtubule binding and motility^55–58^, which is different from our observations with the Kif7-Gli interaction. It is noteworthy, that the Drosophila homolog of Kif7, Costal2 is also proposed to have multiple binding sites for Ci^59, 60^. It is possible that the multi-site interaction may be beneficial for concentrating low copy number transcription factors of the Hh pathway, on microtubules.

Interestingly, our finding of positive feedback regulation implies that by modulating the levels of Kif7 at the cilium tip, Gli acts as a master regulator of its own recruitment to the cilia tip for its activation. Like other transcription factors, Gli is a low copy number and tightly regulated protein, hence the synergistic feedback between Kif7-Gli and Kif7-microtubule interactions is a mechanism that would be advantageous in rapidly concentrating Gli, to the cilium tip upon Hh activation. A similar strategy may be used to concentrate and promote the formation of Gli repressor at the base of the cilium when the Hh pathway is off. Interestingly, even in *Drosophila*, where the Hh pathway is cilium-independent, the Kif7 homolog Costal2 is proposed to tether the Gli homolog Ci to the cytoplasmic microtubules during pathway repression^5^. From an evolutionary perspective, our results indicate that despite the divergence in the utilization of cilium between vertebrates and invertebrates, the fundamental principles by which the microtubule cytoskeleton has been co-opted to regulate Hedgehog signal transduction remains conserved^60^. Our findings also validate that Kif7 and not Kif27 is the mammalian ortholog of fly Costal2 protein that tethers Gli to microtubules.

Gli is the final downstream effector molecule of the Hh signaling pathway and it is therefore a target for regulation of the pathway activity. Aberrant activation of Gli has been reported in multiple human cancers including basal cell carcinomas and glioblastomas through both Hh-dependent and independent mechanisms^12, 61, 62^. An inhibitor that directly targets Gli has therapeutic potential in various cancers with Gli overexpression irrespective of their etiology^63^. Our biochemical analyses of Kif7-Gli interaction and proof-of-principle sequestration experiments suggest that a small coiled-coil peptide from Kif7 can be exploited to sequester Gli in the cytoplasm and restrict its entry into the nucleus **(Fig. 7c).** This strategy could be further developed to downregulate Gli activity as well as a tool for cell biological and optogenetic studies^41–43^. The structural analyses of the Kif7-Gli interaction opens the possibility of a new site on Gli that can be exploited for drug design. On a wider scale, it represents a possible strategy for inhibiting other zinc-finger proteins that play critical roles in gene regulation and may be erroneously activated in disease.

## Methods

### Pull-down assay

Plasmids for the pull-down assay were constructed by cloning human Kif7 and Gli2 sequences into pCMV-BICEP™-4 expression vector followed by truncations using InFusion HD Cloning Kit (Takara). Kif7 fragments were cloned at the MCS1 to express FLAG-tagged Kif7 protein and Gli2 fragments were cloned at the MCS2 to express c-myc-tagged Gli2 protein. Point mutations were made using the Q5® Site-Directed Mutagenesis Kit (NEB). Expi293F^TM c^ells were maintained in Expi293 Expression medium and cultured under 7% CO_2,_ 93% air condition at 37^°C^ with 220rpm shaking. Expi293 cells were transfected with plasmids using ExpiFectamine^TM 2^93 Reagent, grown for 18-22 hours followed by addition of ExpiFectamine^TM 2^93 Transfection Enhancer 1 and grown for another 24 hours. Next the cells were harvested by centrifugation at 2500*xg* for 5min, washed with phosphate buffered saline (PBS) and resuspended in lysis buffer (25mM Tris-HCl, 75mM NaCl, 1mM MgCl_2,_ 0.5mM EDTA, 100μM ATP, 1mM DTT, 0.1% TritonX100 and Halt^TM^ protease inhibitor cocktail; pH=7.5). Cells were lysed by a short sonication and the lysate was cleared by centrifugation at 125000*xg* for 30mins. Anti-FLAG M2 magnetic beads (Sigma Aldrich) pre-equilibrated in the lysis buffer was incubated with the cleared cell lysate (labeled ‘input’) for 1hr at 4°C. DynaMag-Spin was used to collect the supernatant labeled ‘flow through’ and the beads were washed with the buffer 5 times. The last supernatant from the washes was used as the ‘wash’ sample. The final ‘bead’ sample was prepared by resuspending the beads in ∼50ul of buffer. The input, flow through, wash and bead samples were run on SDS-polyacrylamide gel and subjected to Western blotting. Binding was detected using anti-c-myc antibody (Sigma Aldrich) and the expression of the bait-protein was detected from the input sample with anti-FLAG antibody (Sigma Aldrich).

### _P_rotein Expression and Purification

The N-terminal fragment of Kif7 (Uniprot Q2M1P5) (Kif7-DM 1-543aa) was cloned into a pFastBac expression vector (Thermo) that included a tobacco etch virus (TEV) protease cleavable N-terminal 6x His-tag and SUMOstar solubilization tag. To determine which residues to swap in Kif7 460-600aa was used for the coiled-coil. Dimeric Kif7 N-terminal constructs: Kif7-DM (1- 543aa), Kif7-DM_E502A a_nd the Kif7-Kif27 chimeric constructs were expressed in SF9 insect cell line using the Bac-to-Bac® Baculovirus Expressions System (Thermo) with cells grown in HyClone CCM3 SFM (GE Life Sciences) and expressed from P3 virus for 72H at 27°C. Monomeric Kif7 constructs (Kif7-MM 1-398aa) containing the motor domain was cloned into a modified pET-21-a expression vector that contained a TEV protease cleavable N terminal 6x His-tag. Kif7 monomers were expressed in BL21 (DE3) Rosetta (Millipore) E. coli at 18°C with 0.25mM IPTG for 18-20H. Dimeric and monomeric kinesin pellets containing the motor domain of Kif7 were lysed by short sonication in buffer A (50mM phosphate pH 8.0 300mM NaCl 5% glycerol, 1mM MgCl_2 a_nd 25mM imidazole) supplemented with 0.15% tween, 0.5% Igepal, 100μM ATP 2mM TCEP, 1mM PMSF, 75 U benzonase and 1X HALT (Thermo). Lysate was cleared by ultracentrifugation and supernatant was incubated with Ni-NTA for 1H. Resin was washed with buffer A supplemented with 20μM ATP and 0.5mM TCEP and eluted with 400mM imidazole with 100μM ATP. Peak fractions were pooled and, if needed, cleaved overnight by TEV (1/30 w/w) at 4°C. Proteins were further purified by size exclusion chromatography (Superdex 200 10/300GL) in 50mM HEPES pH 7.4 300mM NaCl 5% glycerol 5mM ß-Me 1mM MgCl_2 a_nd 100μM ATP and frozen in liquid nitrogen. Dimerization of a given construct was determinable by gel filtration profiles.

The coiled-coil dimerization domain fragment of Kif7, (Kif7-CC 460-600aa) was cloned into a modified pGEX vector that contained a human rhinovirus (HRV) protease cleavable N terminal GST-tag. The point mutants of Kif7-CC were generated by site-directed mutagenesis using In- Fusion HD cloning kit (Takara). The Kif7 SCC construct (aa488-540) was ordered as a gene- fragment (Genscript) and cloned into pETDuet-1 by In-Fusion HD Cloning Kit. All Kif7-CCs and Kif7-SCC were expressed in BL21 (DE3) Rosetta (Millipore) E. coli at 18°C with 0.25mM IPTG for 18-20hr. The cell pellets were lysed by short sonication in buffer B (PBS pH 7.3, 5% glycerol) supplemented with .15% tween, 0.5% Igepal, 100μM ATP, 2mM TCEP, 1mM PMSF, 75 U benzonase and 1X HALT (Thermo). Lysate was cleared by ultracentrifugation and supernatant was incubated with GST-4B beads (GE Life Sciences) for 1H. Resin was washed with buffer B supplemented with 0.5mM TCEP and eluted with 10mM reduced glutathione in 40mM PIPES pH 7.3 150mM NaCl. Peak fractions were pooled and, if needed, cleaved overnight by GST-HRV-3C (1/40 w/w) at 4°C. Proteins were further purified by size exclusion chromatography (Superdex 200 10/300GL) in 40mM PIPES pH 7.3 150mM NaCl 5% glycerol 2mM TCEP and frozen in liquid nitrogen.

The zinc-finger domain of Gli2 (Uniprot P10070) (Gli2-ZF 418-604aa) was cloned into a modified pET-Duet-1 expression vector that contained an HRV protease cleavable N terminal 6x His-tag. Truncations were made using the InFusion HD Cloning Kit (Takara). The same construct was cloned into a modified pET-Duet-1 vector that contained an HRV protease cleavable N terminal 6x His-tag, and a C-terminal SNAP/GFP tag. Gli2-ZF protein was expressed in BL21 (DE3) Rosetta (Millipore) E. coli at 18°C with 0.25mM IPTG for 18-20hr. Gli2-ZF pellets were lysed by short sonication in buffer C (40mM PIPES pH 7.3 150mM NaCl) supplemented with .15% tween, 0.5% Igepal, 2mM TCEP, 1mM PMSF, 300 U benzonase and 1X HALT (Thermo). Lysate was cleared by ultracentrifugation and supernatant was incubated with Ni-NTA for 1H. Resin was washed with buffer A supplemented with 0.5mM TCEP and eluted with 400mM imidazole. Peak fractions were pooled and, if needed, cleaved overnight by GST-HRV-3C (1/40 w/w) at 4°C. Proteins were further purified by ion-exchange chromatography using a Heparin column; Gli2-ZF proteins are DNA binding and generally elute from the heparin column around 1M NaCl. Proteins were further purified by size exclusion chromatography (Superdex 75 10/300GL and Superdex 200 10/300GL) in 40mM PIPES pH 7.3 150mM NaCl 5% glycerol 2mM TCEP and frozen in liquid nitrogen.

All proteins were greater than 95% pure and eluted as a single peak from the size exclusion chromatography column.

### Bio-Layer Interferometry Assays

BLI experiments were performed in an Octet Red 96 instrument (ForteBio). To quantify Kif7-Gli2 binding, Experiments were performed in an assay buffer containing 40mM TRIS pH7.3, 40mM KCl, 40μM ATP 1mM MgCl_2,_ 0.12% tween and 0.5mM TCEP. In the simplest assay, each dimeric GST-Kif7-CC protein was immobilized on an anti-GST biosensor chip at a concentration of 120nM. Unbound Kif7 was washed away with buffer, and the chips were then dipped into buffer with a range of Gli2-ZF concentrations from 0-1μM to allow for association of the two proteins.

After 300 seconds, they were then moved back into a well with just buffer to allow dissociation for 400 seconds. The equilibrium response of each sensor from three independent repeats at each Gli2 concentration were normalized and plotted against Gli2 concentration and fitted to the following Hill function: Y=(Bmax*X^h^)/(Kd^h^ + X^h^); where B_max i_s the maximum specific binding, K_d i_s the ligand concentration needed to achieve a half-maximum binding at equilibrium and h is the Hill slope.

The DNA competition experiments were performed in the same way with one notable exception: 125μM TRE-2S DNA (a Gli2 DNA binding target) was held constant along the Gli2 titration from 0-1μM. The assays were also performed with HIS-tagged Kif7-DM, Kif7-DM_E502A a_nd the Kif7- Kif27 chimeric proteins, requiring the use of an anti-penta-HIS biosensor chip, and referencing sensors to account for non-specific binding of Gli2 to the HIS sensor. A flipped assay, using HIS-tagged Gli2 on the sensor and a range of Kif7 concentrations in buffer, was also conducted.

### Kif7-Gli2 Complex Stoichiometry Determination

The Kif7-CC and Kif7-DM protein samples were not amenable to dynamic light scattering analysis. To determine the stoichiometry of the Kif7-Gli2 complexes, we purified GFP-tagged constructs of Kif7-DM, Kif7-CC, and Gli2-ZF. Next, the Kif7 (Kif7-CC or Kif7-DM) and Gli2 were mixed at high concentration with Gli2-ZF in molar excess (125μM Kif7 and 750μM Gli2- ZF) and incubated for 15 minutes on ice. After incubation, 200μL of this mix was injected into a Superdex 200 10/300 GL column. Fractions that contained complexed protein were determined by elution volume shift in the gel filtration chromatogram. Complex containing fractions were heated at 50°C for 5 mins and run in SDS-PAGE and quantified in two ways: (1) Scanned the unstained SDS-PAGE gel at 488nm^64^. Since GFP is stable in SDS up to at least 0.5% w/v^65^ ^a^nd SDS-PAGE uses 0.1%, the GFP fluorescence can be quantitatively analyzed on a regular SDS-PAGE gel with this sample preparation protocol^64^. The ratio of GFP fluorescence corresponding to the Kif7 and Gli bands was measured using an Amersham™ Typhoon Scanner and the intensity analyses was performed using ImageJ to obtain the stoichiometry. (2) Quantitative western blot analysis of the complex was performed using a directly labeled 488nM anti-GFP antibody (Santa Cruz). The fluorescence intensity at 488 nm was quantified using an Amersham™ Typhoon Scanner and analyzed with ImageJ. The results from these two independent experiments were consistent with the stoichiometries reported in this study.

### Structural modeling

Primary sequences of the zinc-finger domain of human Gli2 (UniprotKB P10070) (Gli2-ZF, 437- 589aa), the coiled-coil domain of human Kif7 (UniprotKB Q2M1P5) (Kif7-SCC, 481-543aa), the coiled-coil domain of human Kif27 (UniprotKB Q86VH2) (Kif27-CC, 475-570aa) and the N-terminal motor domain of human Kif27 (UniprotKB Q86VH2) (Kif27M, 1-475aa) were retrieved from the database in FASTA format. A preliminary search for homologs to serve as templates for modeling was performed with BLAST^66^ ^a^nd HHBlits^67^ ^a^gainst the SWISS-MODEL template library (SMTL). The following templates with the highest quality predictions (based on target-template alignment) were selected for homology modeling: (i) Gli1 zinc finger (PDB: 2GLI) for Gli2-ZF (ii) ROCK1 coiled-coil (PDB: 3O0Z) for Kif7-CC and Kif27-CC (iii) Kif7 motor domain (PDB: 6MLQ) for Kif27M. The 3D structure of these protein domains was built using ProMod3 in the SWISS-MODEL server^68, 69^. The global and per-residue model quality was assessed using GMQE and QMEAN scoring function^70^. The structural models were visualized in UCSF Chimera^71^. Electrostatic potential of the surfaces were generated from the APBS server^72^.

For obtaining the structural model of the Kif7-SCC:Gli2-ZF protein complex, the Gli2-ZF model was docked on the Kif7-CC model using an automated server, ClusPro 2.0^73^, in which receptor was Kif7-SCC and Gli2-ZF was used as a ligand. Based on different desolvation and electrostatic potential, ClusPro 2.0 can differentiate thousands of conformations of the protein. The generated conformations were further categorized through clustering and 8 most fit structures (which are found to be closest to native structure from X-ray crystallography results) were considered. The interface areas in the structural model of the protein complex were calculated by the PDBsum webserver^74^. These contact residues were mutated to Ala for binding analysis to assess the predicted models.

### Cryo-EM Sample Preparation

For grid preparation, all Kif7-Gli2 preparations were diluted using BRB80 (80 mM 1,4- piperazinediethanesulfonic acid [PIPES], pH 6.8, 1 mM MgCl_2,_ 1 mM ethylene glycol tetraacetic acid [EGTA]). Porcine brain microtubules were prepared from frozen aliquots of 5mg/ml tubulin (Cytoskeleton, Denver, CO) in polymerization buffer (BRB80; 80 mM PIPES, pH 6.8, 1 mM EGTA, 2 mM MgCl_2,_ 3mM GMP-CPP, 10% dimethyl sulfoxide (DMSO)) at 37° C for 2-3 hours. The polymerized microtubules were then incubated at room temperature for several hours or overnight before use. The following day or several hours later the microtubules were spun down on a bench top centrifuge 14,00 RPM 10min to pellet. The pellet was resuspended in polymerization buffer without DMSO. The Kif7 dimer and monomer preparations were made by mixing the Kif7 with the Gli2 or Gli2-SNAP for 1 hour prior to incubation with the microtubules. Kif7-dimer (0.36 mg/ml) plus Gli2 (0.2mg/ml) in BRB plus 2mM ADP, 2mM MgCl_2,_ Kif7-monomer (0.45mg/mL) or Kif7 dimer plus Gli2_SNAP (0.30mg/mL) plus BRB80 with 1mM AMPPNP and 1mM MgCl_2._

All microtubule samples were prepared on 1.2/1.3 400-mesh grids (Electron Microscopy Services) Grids were glow-discharged before sample application. The cryo-samples were prepared using a manual plunger, which was placed in a homemade humidity chamber that varied between 80 and 90% relative humidity. A 4-μl amount of the microtubules at ∼0.5 μM in BRB80 was allowed to absorb for 1 min, and then 4 μl of the different Kif7-dimer or monomer plus Gli2 with and without SNAP were incubated with microtubules on the grid. After a short incubation of 2 min, the sample was blotted (from the back side of the grid) and plunged into liquid ethane.

### EM image acquisition and data processing

Images of frozen-hydrated Kif7-microtubule complexes (see Extended Data Table 1) were collected on a Titan Krios (FEI, Hillsboro, OR) operating at 300 keV or an Arctica (FEI, Hillsboro, OR) equipped with a K2 Summit direct electron detector (Gatan, Pleasanton, CA). The data were acquired using the Leginon automated data acquisition^75, 76^. Image processing was performed within the Appion processing environment^76, 77^. Movies were collected at a nominal magnification of 29000× and 36000x with a physical pixel size of 1.03 and 1.15 Å/pixel respectively. Movies were acquired using a dose rate of ∼4.2 and 5.6 electrons/pixel/second over 9 and 8 seconds yielding a cumulative dose of ∼36 and ∼34 electrons/Å^2^ ^(^respectively). The MotionCor frame alignment program^78, 79^ ^w^as used to motion-correct. Aligned images were used for CTF determination using CTFFIND4^80^ ^a^nd only micrographs yielding CC estimates better than 0.5 at

4 Å resolution were kept. Microtubule segments were manually selected, and overlapping segments were extracted with a spacing of 80 Å along the filament. Binned boxed segments (2.05Å/pixel, 240-pixel box size for the Krios data and 2.30Å/pixel, 192 pixel box size for the Arctica data) were then subjected to reference-free 2D classification using multivariate statistical analysis (MSA) and multi-reference alignment (MRA)^79, 81^. Particles in classes that did not clearly show an 80Å layer line were excluded from further processing.

### Cryo-EM 3D reconstruction

Undecorated 13,14- and 15-protofilament microtubule densities^82^ ^w^ere used as initial models for all preliminary reconstructions. We used the IHRSR procedure^83^ ^f^or multimodel projection matching of microtubule specimens with various numbers of protofilaments^84^, using libraries from the EMAN2 image processing package^85^. After each round of projection matching, an asymmetric backprojection is generated of aligned segments, and the helical parameters (rise and twist) describing the monomeric tubulin lattice are calculated. These helical parameters are used to generate and average 13, 14 and 15 symmetry-related copies of the asymmetric reconstruction, and the resulting models were used for projection matching during the next round of refinement. The number of particles falling into the different helical families varied. Helical families that had enough segments were further refined. Final refinement of microtubule segment alignment parameters was performed in FREALIGN ^86^ ^w^ithout further refinement of helical parameters. The data is unbinned during refinement. The boxsize of the Krios data was binned from 480 to 380 reducing the pixel size from 1.03 to 1.3 Å/pixel. FSC curves were used to estimate the resolution of each reconstruction, using a cutoff of 0.143. To better display the high-resolution features of the 3D map shown, we applied a B-factor of 150 Å, using the program bfactor (http://grigoriefflab.janelia.org).

### Model building in the EM map

The EM-derived structure of AMPPNP-Kif7 bound to microtubules (PDB: 6 MLR, EMD: 9141) was fit as a rigid body into the density for AMPPNP-Kif7 dimer bound to microtubules in the presence of Gli2-SNAP, using the ‘Fit in Map’ utility in UCSF Chimera ^71^. To locate the density corresponding to the Gli2 protein, cryo-EM maps for ADP-Kif7 dimer with Gli2, AMPPNP-Kif7 monomer with Gli2-SNAP and AMPPNP-Kif7 dimer with Gli2-SNAP were superposed via the Kif7 motor domains, and the minimal density common to all three maps was identified. To construct a structural model for Gli2 to fit the density, individual zinc fingers were first extracted from the crystal structure of a five-finger Gli1 in complex with DNA (PDB:2GLI). Next, various combinations of zinc fingers were fit into the minimal density corresponding to Gli2 and the best model was identified through its model-map correlation coefficient and visual inspection. Since the density corresponding to each zinc finger could be fit by any one of the five zinc-fingers of Gli2, the partial structure for Gli2 was modeled as a Poly-Ala sequence. Finally, flexible fitting for the structure comprising of Gli2 zinc fingers, AMPPNP-bound Kif7 motor and α-β tubulin heterodimer was performed with phenix.real_space_refine^87^. The final structure has a model-map correlation of 0.84, bond length RMSD of 0.008 Å and bond angle RMSD of 0.98°. There are no outliers in the Ramachandran map. Cryo-EM reconstructions and their associated models were superposed using the UCSF Chimera “Fit in map” tool^71^.

### Total Internal Reflection Fluorescence Microscopy Assays

*In vitro* TIRF-based microscopy experiments were carried out as described in Subramanian et al., 2013^88^. Microscope chambers were constructed using a 24 x 60 mm PEG-Biotin coated glass slide and 18 x 18 mm PEG coated glass slide separated by double-sided tape to create two channels for exchange of solutions. Standard assay buffer was 1 x BRB80 (80 mM KCl-PIPES at pH 6.8, 2mM MgCl_2 a_nd 1mM EGTA), 1mMATP, 1mMGTP, 0.1%methylcellulose and 3%sucrose. For all experiments, oxygen scavenging mix (OS) comprising of 25 mM glucose, 40 mg/ml glucose oxidase, 35 mg/ml catalase and 0.5% beta-mercaptoethanol was included in the final buffer. Images were acquired using NIS-Elements (Nikon) and analyzed using ImageJ.

Microtubule binding assay: X-rhodamine/HiLyte 647 (1:10 labeled to unlabeled) and biotin (1:10 labeled to unlabeled) labeled microtubules were polymerized in the presence of GMPCPP, a non- hydrolysable GTP-analogue, and immobilized on a neutravidin coated glass coverslip. Coverslips were briefly incubated with casein to block non-specific surface binding before addition of 100nM kinesin and/or varying concentrations of Gli2-ZF in assay buffer and antifade reagent (25mMglucose, 40 mg/ml glucose oxidase, 35 mg/ml catalase and 0.5% β-mercaptoethanol). Images were acquired from multiple fields of the same chamber.

Single molecule assays: Initial fluorescence intensity analysis to determine the oligomerization state of the GFP tagged proteins was performed by immobilizing GFP-tagged protein non- specifically to a glass coverslip. After washing away unbound protein, BRB80-DTT buffer supplemented with OS mix was added to the chamber and images were acquired at 0.37 s intervals on an ANDOR iXon Ultra EMCCD camera with EM gain set to 287. Fluorescent spots were selected by visual inspection and the average intensity of the first five frames was used to compute the initial intensity after background subtraction.

Interaction of single Kif7-DM-GFP molecules with microtubules in the presence of Gli2-ZF was analyzed by immobilizing biotinylated microtubules to a glass coverslip as described earlier, followed by addition of Kif7-DM-GFP (1 nM or 0.1 nM) with increasing concentrations of Gli2- ZF (0 nM, 25 nM, 50 nM) and images were acquired at 0.37 s intervals on an ANDOR iXon Ultra EMCCD camera with EM gain set to 287. The density of tracks in the 1 nM Kif7-DM-GFP data set were too high to perform tracking analysis, so we repeated the experiment with 0.1 nM Kif7- DM-GFP and used that data for analyses.

### Co-sedimentation of Kif7 on GDP-Taxol Stabilized Microtubules

GDP-taxol stabilized microtubules were prepared from purified pre-cleared bovine tubulin as described in Subramanian et al., 2010^89^. Kif7-DM-GFP at 1 µM concentration were incubated with microtubules (0 – 10 µM) in the presence of 1mM ATP, 2mM MgCl_2,_ with or without 5µM Gli for 15 minutes at room temperature in 1x BRB80 supplemented with 0.1 mg/mL BSA and then subjected to sedimentation by ultracentrifugation at 50,000 rpm for 10 minutes at 27°C in a TLA 120.1 rotor (Beckman Coulter). Pellets were subsequently resuspended and the amount of protein in each pellet and supernatant was analyzed by SDS-PAGE. Proteins were stained using Gel Code^TM B^lue Stain (Thermo Fischer Scientific) and scanned using GE Typhoon FLA 9000 Gel Scanner. Dissociation constants were calculated by fitting of the experimental data to a Hill equation using GraphPad Prism.

### Kif7 and Gli2 antibody generation and labeling

The coiled-coil dimerization domain fragment of human Kif7, (Kif7-CC 460-600aa) expressed and purified for the biochemistry experiments was used to produce antibodies in rabbits from Covance Inc. (Princeton, NJ). The Kif7 antiserum was affinity-purified using antigen conjugated CNBr-Sepharose resins. The cDNA sequence encoding 1053-1264 aa mouse was amplified and cloned upstream of sequences encoding a polyhistidine stretch (His) tag in pET23b (Novagen). The mGli2-His recombinant protein was expressed in BL21 (DE3) pLysS cells (Stratagene) and purified under denaturing condition (8 M urea) using the TALON metal affinity resin from Clontech. Purified fusion proteins were used to produce antibodies in guinea pigs from Covance Inc. (Princeton, NJ). The Gli2 antiserum was affinity-purified using antigen conjugated CNBr-Sepharose resins. The Kif7 and Gli2 antibodies were further ‘cleaned’ by incubation with fixed Kif7-/- and Gli2-/- MEFs respectively to reduce non-specific staining in immunofluorescence experiments and labeled with Alex Fluor 647 dye using Molecular Probes kit.

### Cell culture, transfections, and immunofluorescence

HeLa cells were obtained from Robert Kingston (Massachusetts General Hospital). COS7 and NIH3T3 cells were purchased from ATCC and BPS Biosciences respectively. MEFs (WT, Gli2-/-, Gli2-/-Gli3-/-, Kif7-/-) were obtained from Kathryn Anderson (Sloan Kettering Institute), Robert Lipinski (University of Wisconsin), Adrian Salic (Harvard Medical School), and Stephane Angers (University of Toronto). HeLa, NIH3T3 and MEF cells were maintained in high-glucose Dulbecco’s modified Eagle’s medium (DMEM) supplemented with 10% fetal bovine serum (FBS), sodium pyruvate (1mM) and L-glutamine (2 mM). Cells were cultured on clean coverslips under the following conditions: 5% CO_2,_ 95% air condition at 37^°C^.

HeLa and COS7 cells were transfected with plasmids using jetPrime transfection reagent and incubated for 18-22 hours. Next the cells were fixed using a mixture of methanol and acetone (1:1 in volume) for 10 minutes in -20C, washed with washing buffer (PBS+0.05% Tween 20) and mounted on 25mmX75mm glass slides with ProLong Diamond Antifade Mountant with DAPI (Thermo Fisher). Z-stacks were acquired on an inverted Nikon confocal microscope using laser illumination source (405nm, 488nm and 561nm channels corresponding to DAPI, Gli2-Neongreen and Kif7 respectively) with a pinhole of 0.5.

For experiments with NIH3T3, cells were grown to ∼70 confluency followed by transfection with plasmids using jetPrime transfection reagent and incubated for 18-22 hours. For experiments with MEFs, cells were grown to complete confluency. Then both cell types were serum-starved with 0.2% FBS in DMEM to induce ciliogenesis for 24 hours, followed by treatment with 500nM SAG for 12-18 hours before being fixed for immunofluorescence. Cells were fixed using a mixture of methanol and acetone (1:1 in volume) for 10 minutes in -20C, washed with washing buffer (PBS+0.05% Tween 20) for 3 times, blocked with blocking buffer (PBS + 2%BSA; OmniPur BSA; EMD Millipore) for one hour at room temperature. Samples were probed overnight at 4C with the following specific primary antibodies (diluted in blocking buffer): Alexa Fluor 647nm labeled Kif7 antibody (1:10) (in-house), Alexa Fluor 647nm labeled Gli2 antibody (1:10) (in-house), Cyanine3 or Alexa-488 labeled polyclonal γ-tubulin antibody (1:1000) (Thermo Fisher) and Alexa Fluo 488nm labeled acetylated α-tubulin antibody (1:500) (Santa Cruz Biotechnology). Samples were mounted on a 25mmX75mm glass with ProLong Diamond Antifade Mountant (Thermo Fisher). Z-stacks were acquired on an inverted Nikon confocal microscope using laser illumination sourc.3e (488nm, 561nm and 647nm channels respectively) with a pinhole of 1. Z-projections were generated by sum of images from planes that included the entire cilium.

### Quantification and Statistical Analysis

ImageJ was used to assess GFP/Alexa 647 fluorescence intensities on microtubules. For all average intensity per pixel values recorded, a rectangular area along each microtubule was selected with a width of 3 pixels. Background intensities were also subtracted locally from regions of interest of the same area around the selected microtubule. Intensities were not analyzed for microtubules found at the edges of the camera’s field of view.

For single molecule residence time analysis, fluorescent spots along each microtubule were analyzed using the ImageJ tracking plugin TrackMate to measure the track durations of the single kinesin molecules. The cumulative frequency plots of residence time were fitted to the following bi-phasic association function model in GraphPad Prism:

Span^Fast=^(Plateau-Y0)*Percent^Fast*^.01 Span^Slow=^(Plateau-Y0)*(100-Percent^Fast)^*.01

Y=Y0+ Span^Fast*^(1-exp(-K^Fast*^X)) + Span^Slow*^(1-exp(-K^Slow*^X))

And Tau^Fast (^reciprocal of K^Fast)^ represents lifetime of single Kif7-DM-GFP molecules and Tau^Slow (^reciprocal of K^Slow)^ are likely to represents lifetime of single Kif7-DM-GFP molecules bound by Gli2-ZF.

The cumulative frequency plot of residence time of Kif7-DM in the absence of Gli2-ZF was not fit since a high fraction of the events in this condition have a very short the residence times of 1-2 frames. This indicates that the true lifetime is lower than the time resolution of our experiment (0.37 s per frame). For the same reason, the Tau^Fast v^alues are likely to represent the upper limit for the lifetime of un-complexed Kif7-DM molecules.

All TIRFM data were analyzed using GraphPad Prism. ‘‘N’’ numbers in experiments refer to the unique number of microtubules or single kinesin molecules (as mentioned in the figure legends) used for the dataset. Details of fit parameters can be found in the corresponding figure legends. ImageJ was used to analyze intensity of protein bands for stoichiometry determination. A rectangular box was drawn around the protein band of interest for intensity readout. Local background was subtracted using a 3-pixel radius around the region of interest.

Octet Data Analysis software was used to extract the binding response for BLI data. The responses were plotted as a function of protein concentration and fit to Hill curves using GraphPad Prism. The binding responses were normalized between 0 and 100 prior to curve fitting.

For structure fitting, atomic models were fit into the cryo-EM density using UCSF Chimera and model to map correlation coefficients were calculated using the ‘Fit in Map’ utility.

For quantification of Kif7 intensity in the cilia, images of cilia were analyzed on ImageJ. The Alexa647 intensity profile along the length of each cilium was measured from tip to base. The acetylated α-tubulin channel was used as a marker for cilia length and the γ-tubulin channel was used as a marker for the base of the cilia. Intensity profiles of different cilia lengths were aligned by their tips and averaged. The integrated intensity profiles were plotted using GraphPad prism. Statistical details can be found in the results section and corresponding figure legends.

For cytoplasmic sequestration quantification, images of cells were analyzed on ImageJ. The nucleus boundary points (including the nucleolus pattern) were identified by using automatic thresholding on DAPI channel (Otsu method in Image J). The threshold value was obtained from auto-thresholding the brightest frame of confocal z-stacks images. This threshold value was used to create binary masks on individual frame of Z-stacks. These masks created by applying threshold on the DAPI channel (called “DAPI Nucleus mask”) were used to measure Gli2-neongreen intensity in the nucleus. Finally, the nuclear Gli2-neongreen intensity was integrated after background subtraction by rolling ball (r=100 pixels) in image J. Next, to create a mask in the Gli2-neongreen for the whole cell boundary an ImageJ macro was applied that automatically determined the threshold value with the best-match mask in the nucleus area (“Gli2 Nucleus mask”) with ”DAPI Nucleus mask”. The best match was selected by finding the maximum value in XNOR operation between “Gli2 Nucleus mask” and “DAPI Nucleus mask”. With the given threshold value, a binary mask over the whole cell area was created (“Gli2 whole cell mask”) and again the Gli2-neongreen intensity in the whole cell was integrated after background subtraction by rolling ball (r = 100 pixels).

### Data and materials availability

The datasets generated and/or analyzed during the current study are available as supplemental table. Any additional data are available from the corresponding author on reasonable request.

### Code availability

All custom codes used for analyses of the data are available from the corresponding author on reasonable request.

## Acknowledgements

We thank Dr. Robert E. Kingston (Massachusetts General Hospital), Dr. Kathryn V. Anderson (Sloan Kettering Institute), Dr. Robert Lipinski (University of Wisconsin), Dr. Adrian Salic (Harvard Medical School), and Dr. Stephane Angers (University of Toronto) for kindly providing us with the HeLa and MEF (WT, Gli2-/-, Gli2-/-Gli3-/-, Kif7-/-) cell lines. We also thank Dr. Mu He (University of California San Francisco) for valuable discussions. This work was supported by a grant to R.S. from American Cancer Society. R.S. is a Pew Biomedical Scholar and a recipient of the Smith Family Award for Excellence in Biomedical Research.

## Author Contributions

R.S., and F.H. designed the project. R.S., E.M.W.-K and R.A.M. designed the cryo-EM experiments; F.H., performed pull-down assays, homology modelling, in vitro reconstitution, TIRF microscopy, co-sedimentation and cell biological experiments; C.F. purified recombinant proteins and performed stoichiometry and BLI assays; Q.Y. performed initial pull-down assays; E.M.W.-K. performed cryo-EM experiments and obtained 3D reconstructions; N.M. did model building in EM map; P.-I.K. performed cytoplasmic sequestration image analysis; F.H., C.F. and R.S. wrote the manuscript. All authors discussed the results and reviewed the manuscript.

Extended data and Supplementary Information are available for this paper.

