## Supplementary Info for "Cytoskeletal regulation of a transcription factor by DNA mimicry"

### Extended Data Figure 1

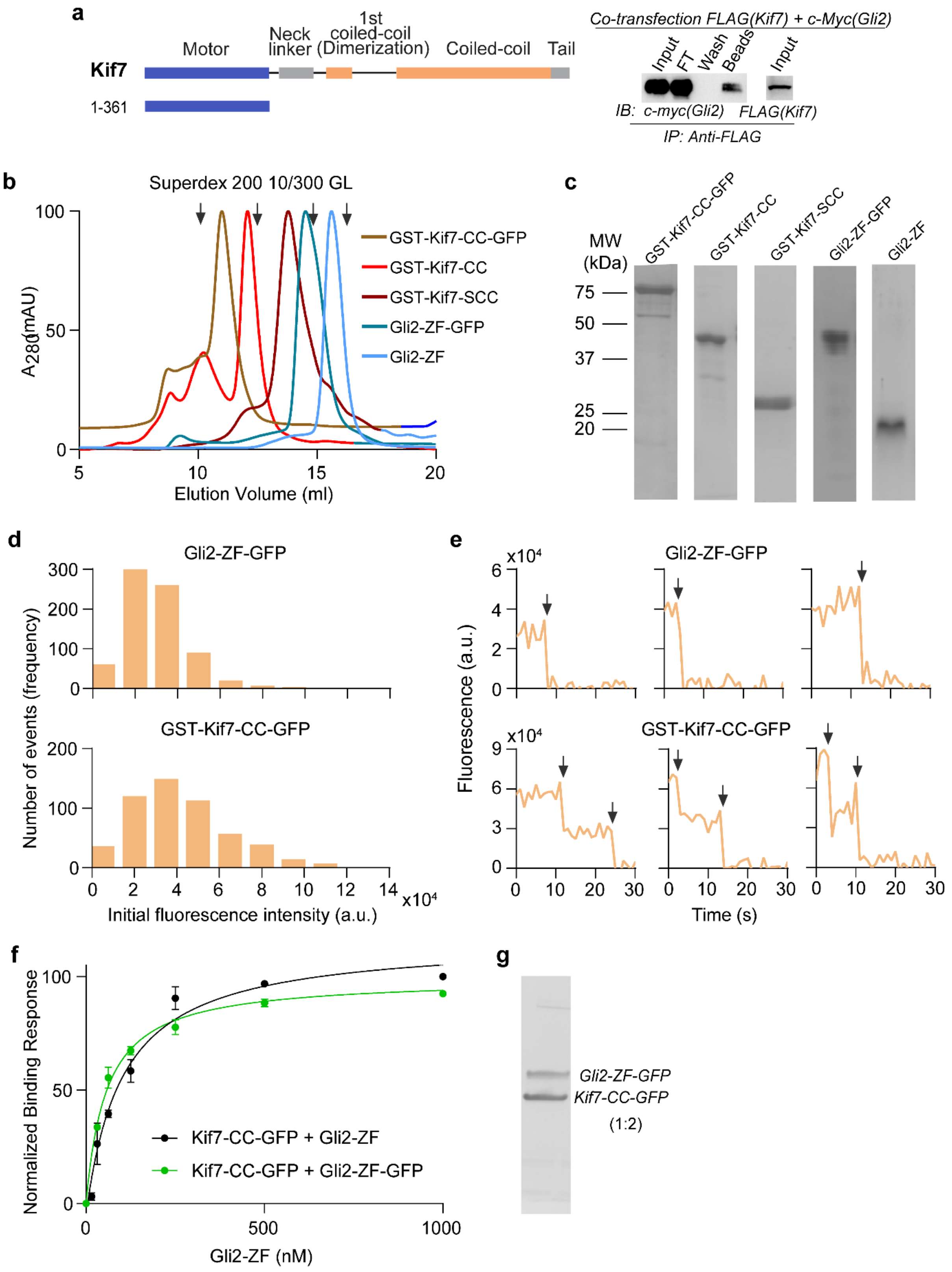

**Extended Data Figure 1 | Purification, stoichiometry analysis and binding of recombinant Kif7-CC and Gli2-ZF proteins. a,** Pull-down of c-Myc-Gli2 (418-594aa) and FLAG-Kif7 (1-361aa) after co-transfection in Expi293F cells. Immunoprecipitation (IP) using anti-FLAG magnetic beads. Input (cell lysate), FT (flow through), wash and beads samples were immunoblotted (IB) with anti c-myc antibody to detect Gli2. Immunoblot with anti-FLAG antibody was used to detect the expression of Kif7. **b,** Chromatograms from size exclusion chromatography of GST-Kif7-CC-GFP, GST-Kif7-CC, GST-Kif7-SCC, Gli2-ZF-GFP and Gli2-ZF on Superdex 200 10/300 GL. Arrows indicate the elution volumes of the following standards: *(left to right)* 1-ferritin (440kDa), 2-aldolase (158kDa), 3-ovalbumin (44kDa) and 4-carbonic anhydrase (29kDa). **c,** SDS-PAGE of purified GST-Kif7-CC-GFP, GST-Kif7-CC, GST-Kif7-SCC, Gli2-ZF-GFP and Gli2-ZF. MW – molecular weight markers. **d,** Single molecule fluorescence intensity histograms of Gli2-ZF-GFP (Intensity = $3.0 \times 10^4 \pm 1.4 \times 10^4$, N = 746) and GST-tagged Kif7-CC-GFP (Intensity = $4.2 \times 10^4 \pm 1.9 \times 10^4$, N = 536). Intensities are reported as mean ± standard deviation. **e,** Single molecule photobleaching traces for Gli2-ZF-GFP ($1^{st}$ row) and GST-Kif7-CC-GFP ($2^{nd}$ row). Background subtracted integrated fluorescence intensity versus time plots used for step photobleaching analysis. Photobleaching steps are indicated by arrows. **f,** BLI assay to quantitatively examine the binding affinity of Kif7-CC-GFP to Gli2-ZF (black) & Gli2-ZF-GFP (green). Data represent mean and standard deviation from three independent repeats. The plots of binding response versus Gli2-ZF concentration were fit to a Hill equation to determine equilibrium dissociation constants ($K_d$). For Kif7-CC-GFP + Gli2-ZF: $K_d = 65 \pm 22$nM, for Kif7-CC-GFP + Gli2-ZF-GFP: $K_d = 57 \pm 9$nM. [Note: GFP-tagged versions of proteins show a consistent increase in $K_d$ that is within error range]. **g,** Western blot of peak complex fraction from size exclusion chromatography of Kif7-CC-GFP and Gli2-ZF-GFP (see Fig. 1d). Stoichiometry of components in complex is indicated in parenthesis.

### Extended Data Figure 2

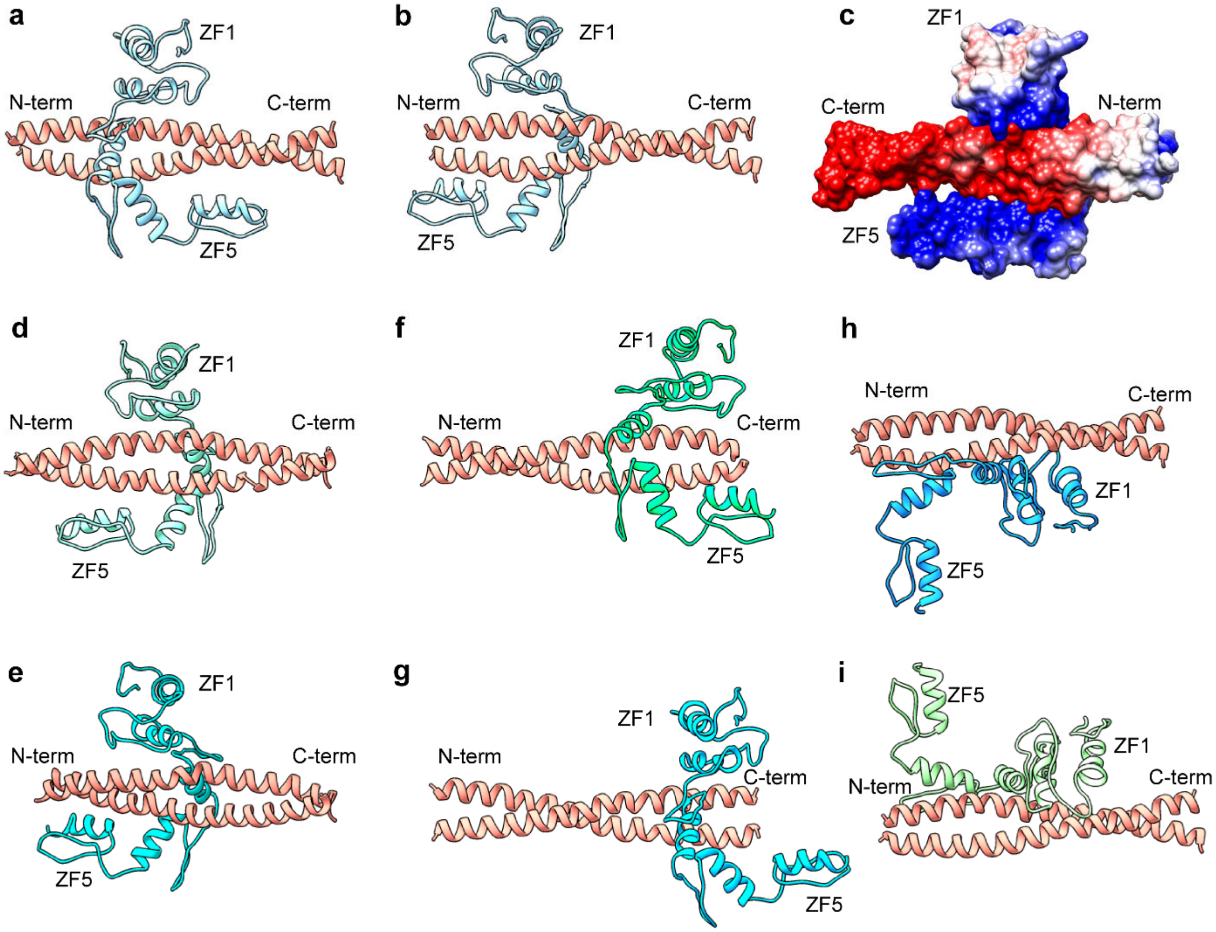


**Extended Data Figure 2 | Structural models of Kif7-CC-Gli2-ZF protein complex.** High confidence structural models of the Kif7-CC-Gli2-ZF protein complex from docking analysis (ClusPro 2.0). Gli2-ZF model is represented in blue and green colored ribbon diagrams and Kif7-CC models represented in salmon colored ribbon diagrams in the different models. **a-b,** Possible structural models that cannot be distinguished from our experiments. **c,** Overall electrostatic surface representation of the complex in **a**. **d-g,** Structural models that are eliminated by mutagenesis experiments (specifically Kif7-CC S2-mut, S3-mut1 and S3-mut2, see Extended Data Fig.3e,f,i). **h-i,** Structural models that are eliminated by the DNA competition assay (See Fig. 2d).

**Extended Data Figure 3**

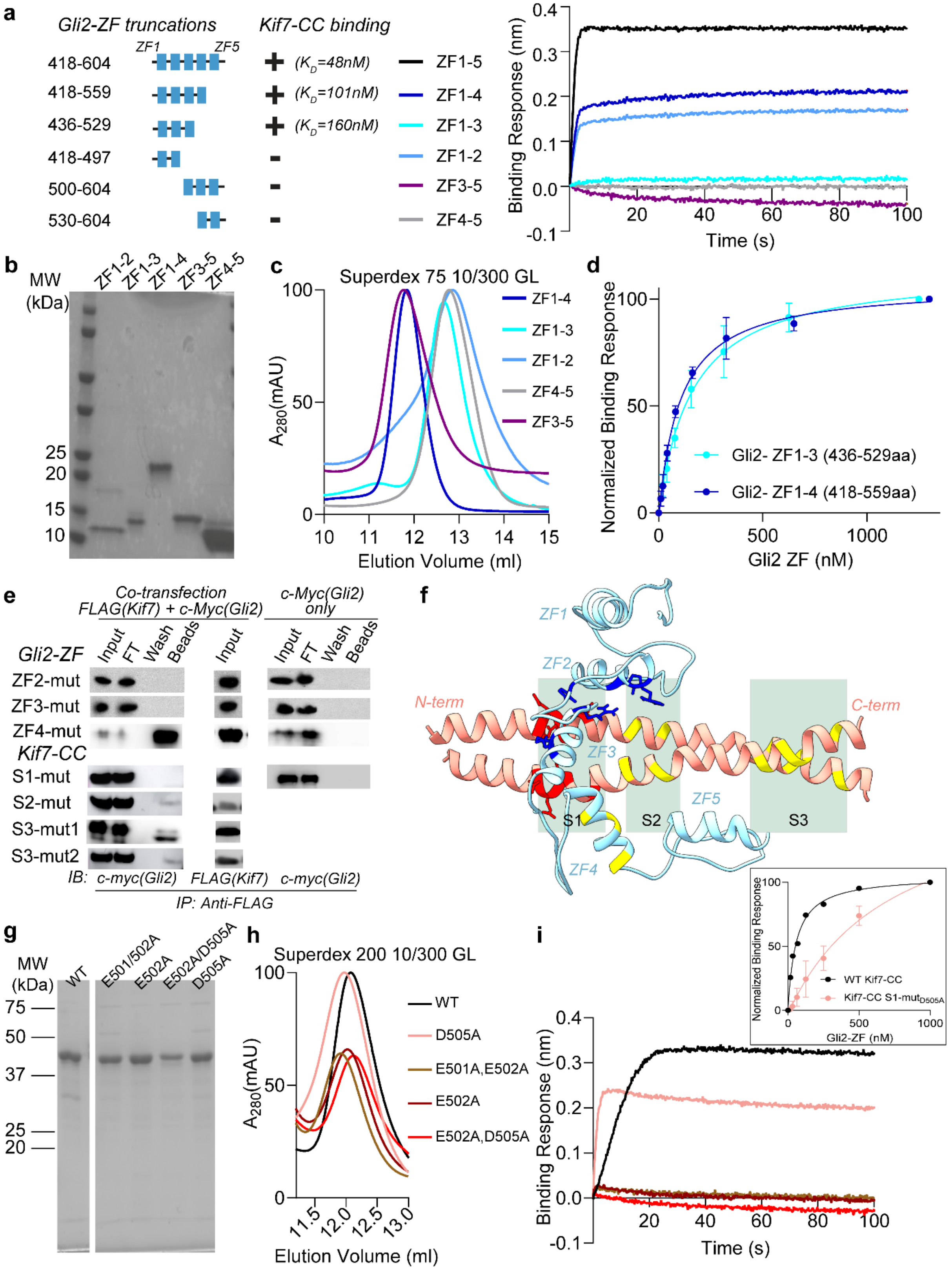

**Extended Data Figure 3 | Site-directed mutagenesis to validate the structural model of the Kif7-CC:Gli2-ZF complex. a,** Schematic of Gli2-ZF truncations and summary of their binding to Kif7-CC *(left panel)*. Raw traces from the association step in the BLI assay performed with soluble 125nM Gli2-ZF truncations and Kif7-CC *(right panel)*. Equilibrium dissociation constants $K_d$ (shown in parenthesis) were determined by fitting complete binding curves to Hill equation (Extended Data Fig. 3d). Data is representative trace of a minimum of three repeats. **b,** SDS-PAGE of purified Gli2-ZF truncation proteins. MW – molecular weight markers. **c,** Chromatograms from size exclusion chromatography of the Gli2-ZF truncation proteins on Superdex 75 10/300 GL. **d,** Binding response in BLI assay for the binding of Kif7-CC with Gli2 truncation constructs: ZF1-3 and ZF1-4. Data represent mean and standard deviation from three independent repeats. The plots of binding response versus Gli2-ZF concentration were fit to a Hill equation to determine equilibrium dissociation constants ($K_d$). For Gli2-ZF 1-4: $K_d = 103 \pm 15$nM and Gli2-ZF 1-4: $K_d = 160 \pm 40$nM. **e,** Pull-down experiment with c-Myc-Gli2 (418-594aa) and FLAG-Kif7-CC (460-600aa) mutant proteins after co-transfection in Expi293F cells. Immunoprecipitation (IP) using anti-FLAG magnetic beads. Input (cell lysate), FT (flow through), wash and beads samples were immunoblotted (IB) with anti c-Myc antibody to detect Gli2. Cells that were only transfected with c-Myc-Gli2 (418-594aa) were included as a negative control. The blots are representative images of a minimum of three repeats. Gli2-ZF mutants are as follows: ZF2-mut (H493A, R496A), ZF3-mut (H503A, R516A, K521A), ZF4-mut (R550A, K552A, R556A). Kif7-CC mutants are as follows: S1-mut (E500A, E501A, E502A, D505A), S2-mut (E511A, E515A), S3-mut1 (E526A, E529A) and S3-mut2 (E530A, R535A). **f,** Structural model of Kif7-SCC:Gli2-ZF protein complex. In dark blue are residues on Gli2-ZF which when mutated to alanine abolish binding with Kif7-CC. In red are residues on Kif7-CC which when mutated to alanine abolish binding to Gli2-ZF. In yellow are residues on both Kif7-CC and Gli2-ZF which when mutated to alanine did not abolish binding. Sections highlighted in green (S1, S2, S3) represent 3 patches of residues on Kif7-CC that were predicted as potential Gli binding sites by PDBsum analysis. **g,** SDS-PAGE of purified Kif7-CC S1 point mutant proteins. MW – molecular weight markers. **h,** Chromatograms from size exclusion chromatography of the Kif7-CC S1 point mutant proteins on Superdex 200 10/300 GL. **i,** Raw traces from the association step in the BLI assay performed with Kif7-CC S1 point mutant proteins with Gli2-ZF: D505A, E501A/E502A, E502A, E502A/D505A. ***Inset*,** Binding response in BLI assay for D505A Kif7-CC mutant to Gli2-ZF. Data represent mean and standard deviation from three independent repeats.

Extended Data Figure 4

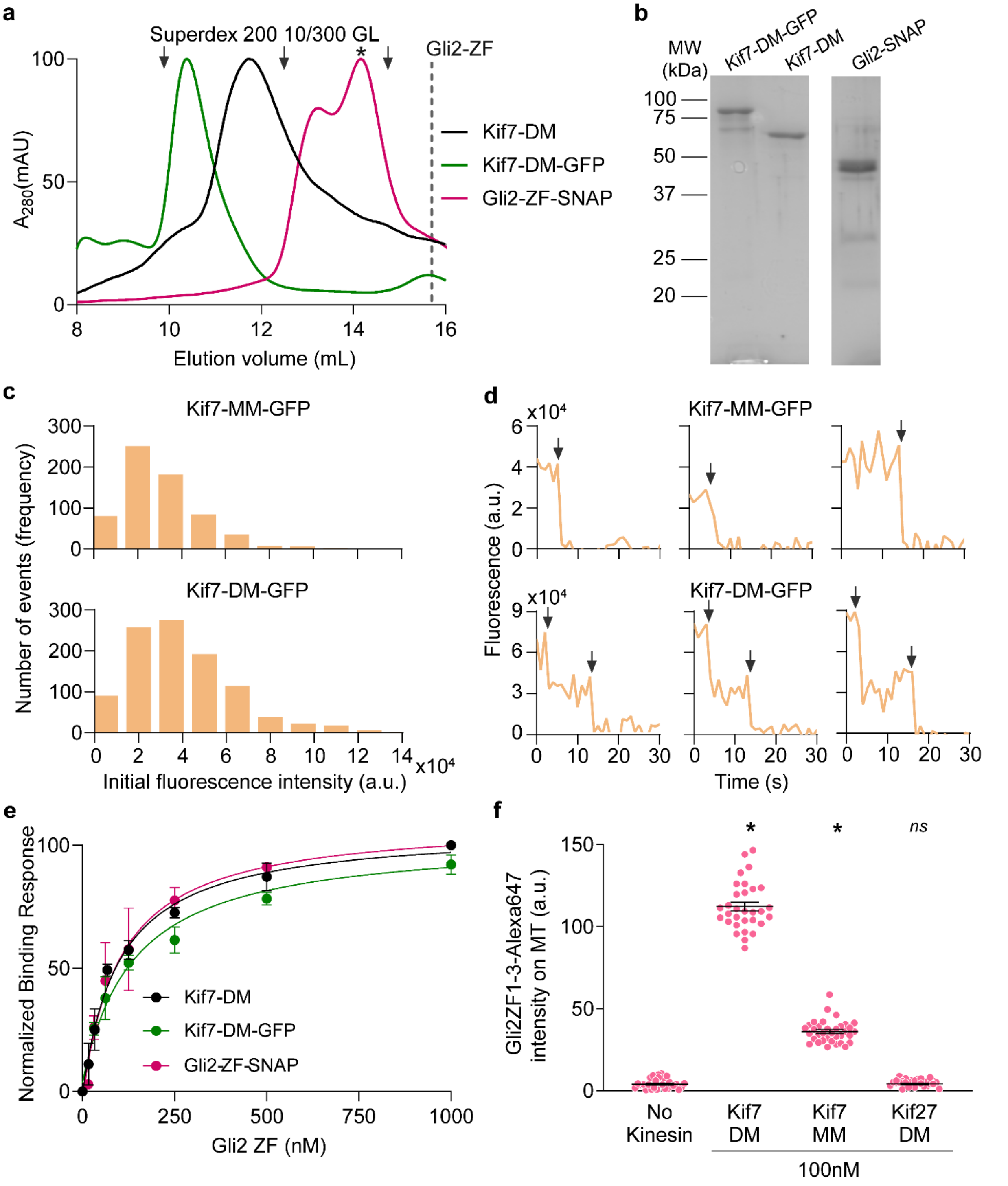


**Extended Data Figure 4 | Purification and stoichiometry analysis of recombinant Kif7-DM Kif7-MM and Gli2-ZF-SNAP proteins. a,** Chromatograms from size exclusion chromatography of Kif7-DM (black), Kif7-DM-GFP (green) and Gli2-ZF-SNAP (magenta, * shows the peak that was used for experiments) on Superdex 200 10/300 GL. Dotted line indicates elution volume of Gli2-ZF on the same column. Arrows indicate the elution volumes of the following standards: *(left to right)* 1-ferritin (440kDa), 2-aldolase (158kDa) and 3-ovalbumin (44kDa). **b,** SDS-PAGE of purified Kif7-DM, Kif7-DM-GFP and Gli2-ZF-SNAP. MW – molecular weight markers. **c,** Fluorescence intensity histograms of Kif7-MM-GFP (Intensity = $3.0 \times 10^4 \pm 1.7 \times 10^4$, N = 648) and Kif7-DM-GFP (Intensity = $4.0 \times 10^4 \pm 2.0 \times 10^4$, N = 1015). Intensities are reported as mean ± standard deviation. **d**, Single molecule photobleaching traces for Kif7-MM-GFP (top) and Kif7-DM-GFP (bottom). Background subtracted integrated fluorescence intensity versus time plots used for step photobleaching analysis. Photobleaching steps are indicated by arrows. **e,** BLI assay to quantitatively examine the binding affinity of Kif7-DM (black) and Kif7-DM-GFP (green) to Gli2-ZF & Kif7-DM to Gli2-ZF-SNAP (maroon). Data represent mean and standard deviation from three independent repeats. The plots of binding response versus Gli2-ZF concentration were fit to a Hill equation to determine equilibrium dissociation constants ($K_d$). For Kif7-DM + Gli2-ZF: $K_d = 99 \pm 25$nM, for Kif7-DM + Gli2-ZF-SNAP: $K_d = 99 \pm 23$nM, for Kif7-DM-GFP + Gli2-ZF: $K_d = 138 \pm 41$nM, for [Note: GFP-tagged versions of proteins show a consistent increase in $K_d$ that is within error range]. **f,** Scatter plot of Gli2-ZF-1-3-Alexa647 intensity per pixel on microtubules (MT) represents recruitment of Gli on MT by Kif7-MM and Kif7-DM. The no kinesin condition was a control for non-specific binding of Gli2-ZF-1-3 to MT and Kif27-DM was included as a negative control for Gli binding. Assay conditions: 100nM Gli2-ZF-1-3-SNAP-Alexa647 with 100nM of kinesin in each case. N $\geq$ 30 microtubules for each condition. One-way ANOVA ($p < 0.0001$) and post-hoc analysis ($*p < 0.0001$ in Dunnett's multiple comparisons test) show statistically significant differences in Gli intensity compared to no kinesin control; *ns* is not significant ($p > 0.5$).

### Extended Data Figure 5

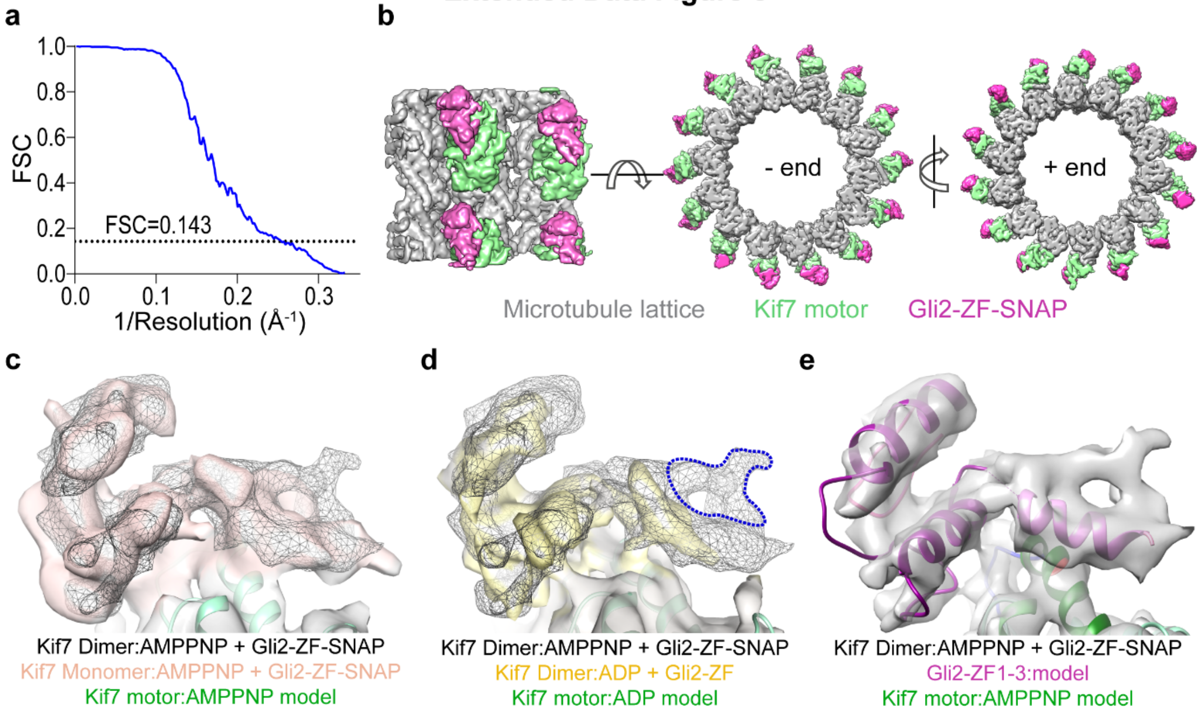


**Extended Data Figure 5 | Cryo-EM maps of Kif7 and Gli2-ZF on microtubules. a,** Fourier shell correlation (FSC) curve for Kif7-DM bound to GMPCPP microtubules in the presence of AMPPNP and Gli2-ZF-SNAP. **b,** Cryo-EM reconstruction of dimeric Kif7 motor domain (green) in complex with AMPPNP bound on the GMPCPP-microtubule lattice (grey) in the presence of Gli2-ZF-SNAP (pink), shown in two orientations and from the –end and +end of the microtubule. Density of Gli2-ZF-SNAP is seen only attached to the Kif7 motor domain and no extra density is seen along the microtubule lattice. **c,** EM reconstructions of AMPPNP-bound Kif7-DM with Gli2-ZF-SNAP (black mesh) and AMPPNP-bound Kif7-MM with Gli2-ZF-SNAP (orange) superposed via the Kif7 motor domain. Structural model for AMPPNP-bound Kif7 motor (green) is shown. **d,** EM reconstructions of AMPPNP-bound Kif7-DM with Gli2-ZF-SNAP (black mesh) and ADP-bound Kif7-DM with Gli2-ZF (yellow) superposed via the Kif7 motor domain. Structural model for AMPPNP-bound Kif7 motor (green) is shown. Blue dotted region indicates density seen only with Gli2-ZF-SNAP and not with Gli2-ZF (thereby corresponding to part of SNAP peptide). **e,** Cryo-EM reconstructions of AMPPNP-bound Kif7-DM with Gli2-ZF-SNAP. Structural models for AMPPNP-bound Kif7 motor (green) and Gli2-ZF consisting of 2 complete and one partial zinc finger (magenta) are shown. Helix $\alpha$-2 (dark green) and loop L2 (blue) of Kif7 make contacts with the density for Gli2-ZF-SNAP. See also Supplementary Video 1.

### Extended Data Figure 6

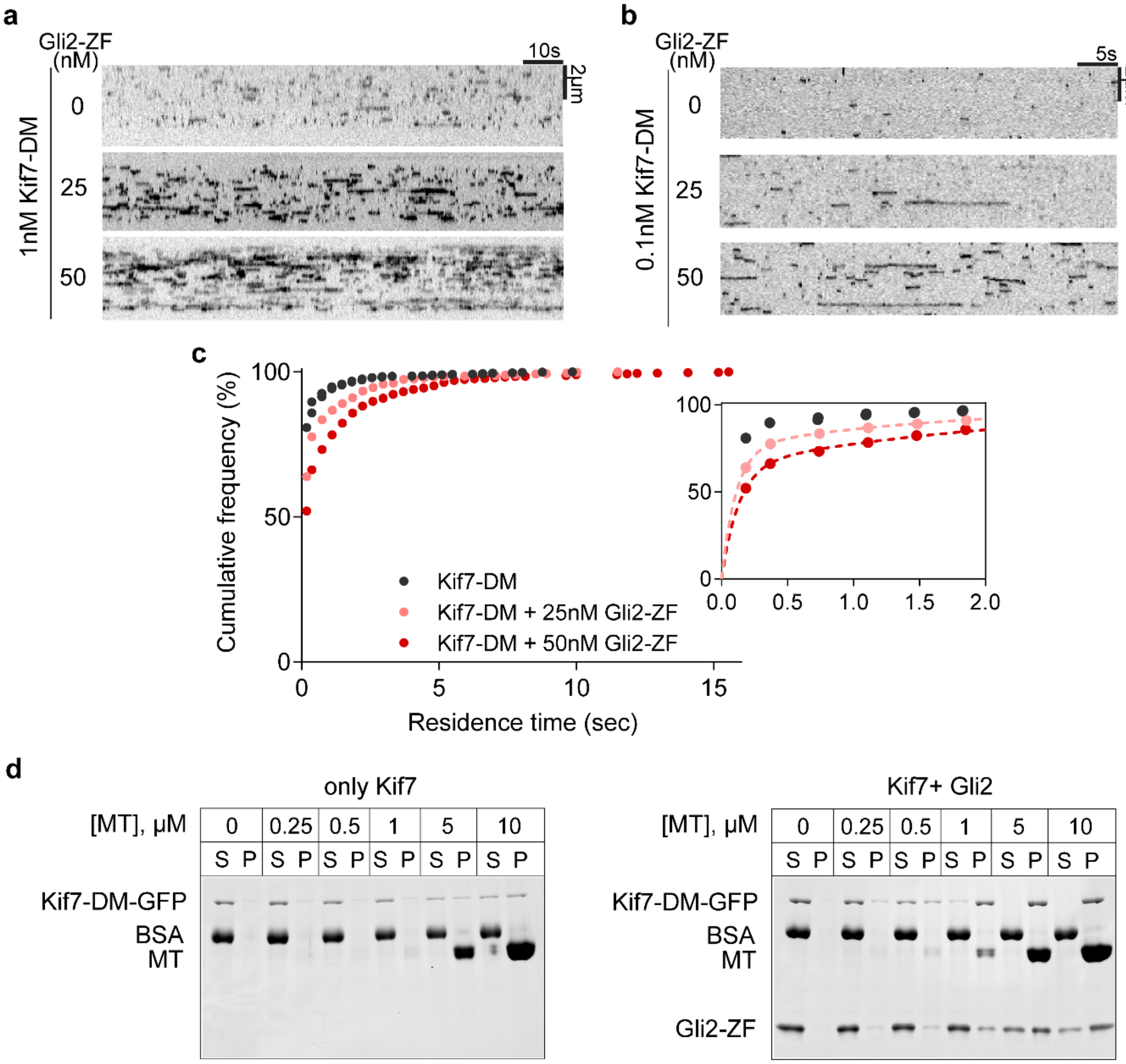


**Extended Data Figure 6 | Single molecule residence time analysis and microtubule co-sedimentation assay with Kif7-DM-GFP in the presence of Gli2-ZF. a,** Representative kymographs from assays to visualize single molecules of Kif7-DM-GFP (1nM) on X-rhodamine labeled microtubules with increasing Gli2-ZF concentrations (0, 25 & 50 nM). **b,** Representative kymographs from assays to visualize single molecules of Kif7-DM-GFP (0.1nM) on X-rhodamine labeled microtubules with increasing Gli2-ZF concentrations (0, 25 & 50 nM). **c,** Cumulative frequency distribution plots of Kif7-DM-GFP residence time on microtubules from analysis of 0.1

nM Kif7-DM-GFP data set with increasing Gli2-ZF concentrations. Numbers of observed events at different Gli2-ZF concentrations: 0 nM, 1191; 25 nM, 576; 50 nM, 817. Inset shows expanded view of the same data. Dotted lines represent bi-phasic association functions fitted to the 25nM and 50nM Gli data sets. [Parameters from fit: (i) For 25 nM Gli; $R^2$ = 0.99; $Tau^{Fast}$ = 0.11 s; $Tau^{Slow}$ = 1.64 s; (ii) For 50 nM Gli; $R^2$ = 0.99; $Tau^{Fast}$ = 0.13 s; $Tau^{Slow}$ = 2.05 s]. See Methods for details. **d,** SDS-PAGE analysis of 1 µM Kif7DM-GFP co-sedimentation on GDP-taxol stabilized microtubules (0 – 10 µM) in the presence of 1 mM ATP, 2mM $MgCl_2$, with or without 5 µM Gli2-ZF and the analysis of 5 µM Gli2-ZF non-specific co-sedimention on microtubules in the absence of Kif7 (S: Supernatant, unbound fraction. P: Pellet, bound fraction. BSA: Bovine Serum Albumin, gel loading control).

### Extended Data Figure 7

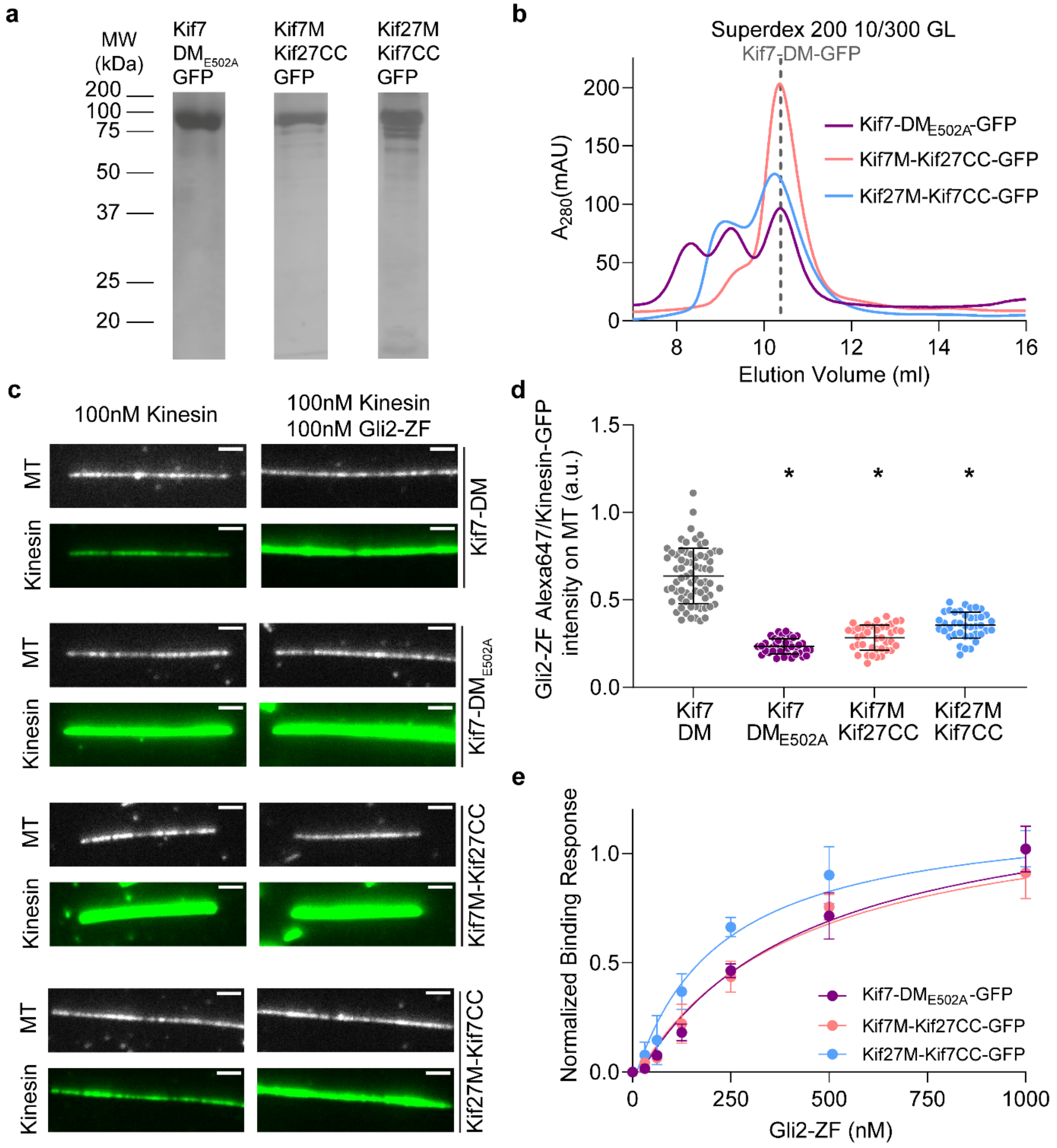


**Extended Data Figure 7 | Purification and binding of recombinant dimeric Kif7 mutant and chimera proteins to Gli2-ZF in solution and on microtubules. a,** SDS-PAGE of purified Kif7-$DM_{E502A}$-GFP, Kif7M-Kif27CC-GFP and Kif27M-Kif7CC-GFP. MW – molecular weight markers. **b,** Chromatograms from size exclusion chromatography of Kif7-$DM_{E502A}$-GFP (purple),

Kif7M-Kif27CC-GFP (red) and Kif27M-Kif7CC-GFP (blue) on Superdex 200 10/300 GL. Dotted line indicates elution volume of Kif7-DM-GFP on the same column. **c,** Representative images of microtubule (MT) and Kinesin-DM-GFP (Kinesin) in the absence (left) or presence of Gli2-ZF (right, 100nM). Fluorescence intensities were scaled similarly for images. Scale bars represent 2μm. **d,** Recruitment of Gli to microtubules (MT) by Kif7-DM-GFP (grey), Kif7-$DM_{E502A}$-GFP (purple), Kif7M-Kif27CC-GFP (red) and Kif27M-Kif7CC-GFP (blue). Scatter plot shows the ratio of Gli2-ZF-Alexa647 intensity to Kinesin GFP intensity on microtubules. Assay conditions: 100nM Gli2-ZF-SNAP-Alexa647 with 100nM of kinesin in each case. $N \geq 40$ for each condition. One-way ANOVA ($p < 0.0001$) and post-hoc analysis ($*p < 0.0001$ in Dunnett's multiple comparisons test) show statistically significant differences in Gli intensity compared to Kif7-DM-GFP control. **e,** BLI assay to quantitatively examine the binding affinity of Kif7-$DM_{E502A}$-GFP (purple), Kif7M-Kif27CC-GFP (red) and Kif27M-Kif7CC-GFP (blue) to Gli2-ZF. Data represent mean and standard deviation from three independent repeats. The plots of binding response versus Gli2-ZF concentration were fit to a Hill equation to determine equilibrium dissociation constants ($K_d$). For Kif7-$DM_{E502A}$-GFP: $K_d = 414 \pm 103$nM, for Kif7M-Kif27CC-GFP: $K_d = 397 \pm 88$nM and for Kif27M-Kif7CC-GFP: $K_d = 217 \pm 70$nM.

### Extended Data Figure 8

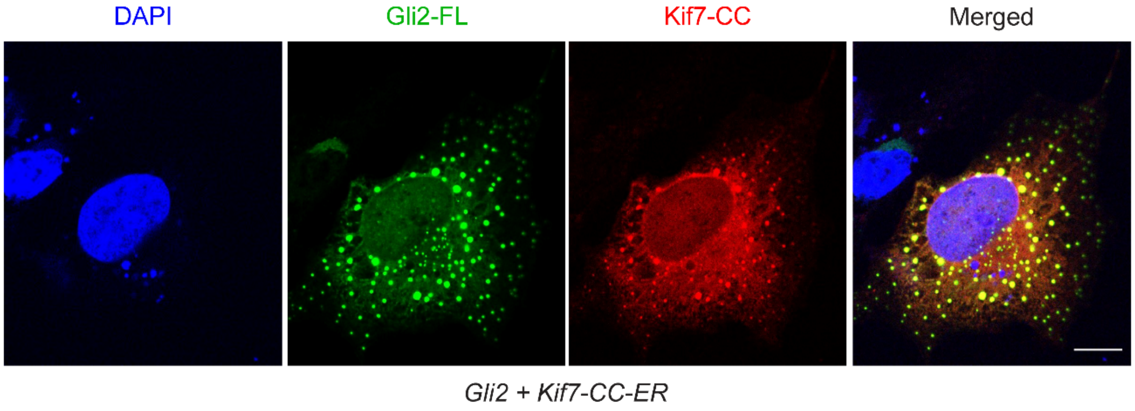


**Extended Data Figure 8 | Sequestration of overexpressed Gli2 using re-engineered DNA-mimicking Kif7-CC in COS7 cells.** Localization of co-transfected mRuby-tagged Kif7-CC (460-600aa) with ER retention tag and mNeonGreen-tagged full length Gli2 in COS7 cells. Kif7-CC shows localization at the endoplasmic reticulum in the cytoplasm. DAPI staining was used to mark the nucleus. Gli2 is sequestered in the cytoplasm and shows co-localization with Kif7-CC both on the endoplasmic reticulum and in cytoplasmic punctae. Scale bars represent 10 µm.

**Extended Data Table 1. Data collection and reconstruction of cryo-EM maps**

| | Ki7-MM-AMPPNP-Gli-SNAP (14 protofilaments) | Ki7-DM-AMPPNP-Gli-SNAP (14 protofilaments) | Kif7-DM-ADP-Gli (15 protofilaments) |
|---|---|---|---|
| **Data collection** | | | |
| Microscope | Titan Krios (FEI) | Titan Krios (FEI) | Arctica (FEI) |
| Voltage (kV) | 300 | 300 | 200 |
| Nominal magnification* | 29,000X | 29,500X | 36,000X |
| Cumulative exposure dose ($e^- Å^{-2}$) | 36 | 36 | 34 |
| Exposure rate ($e^-$/pixel/sec) | 4.2 | 4.2 | 5.6 |
| Detector | K2 Summit | K2 Summit | K2 Summit |
| Pixel size (Å)* | 1.03 | 1.03 | 1.15 |
| Defocus range (μm) | 0.04-4.5 | 0.17-4.8 | 0.04-4.5 |
| Average defocus (μm) | 1.18 | 1.29 | 1.17 |
| Micrographs Used | 687 | 1,364 | 2,199 |
| Total extracted helical segment (no.) | 24,677 | 50,639 | 25,474 |
| Refined helical segment (no.) | 11,956 | 32,251 | 16,880 |
| **Reconstruction** | | | |
| Final helical segments (no.) | 9,952 | 25,730 | 16,880 |
| Symmetry imposed | HP | HP | HP |
| Resolution (global) FSC 0.143 | 4.8 | 3.89 | 4.3 |

**Supplemental Video 1. Cryo-EM reconstructions of AMPPNP-bound Kif7-DM with Gli2-ZF-SNAP.** Structural models for AMPPNP-bound Kif7 motor (green) and Gli2-ZF consisting of 2 complete and one partial zinc finger (magenta) are shown.
